# Divergent microbial traits influence the transformation of living versus dead root inputs to soil carbon

**DOI:** 10.1101/2022.09.02.506384

**Authors:** Noah W. Sokol, Megan M. Foley, Steven J. Blazewicz, Amrita Battacharyya, Katerina Estera-Molina, Mary Firestone, Alex Greenlon, Bruce A. Hungate, Jeffrey Kimbrel, Jose Liquet, Marissa Lafler, Maxwell Marple, Peter Nico, Eric Slessarev, Jennifer Pett-Ridge

## Abstract

Soil microorganisms influence the global carbon cycle by transforming plant inputs into soil organic carbon (SOC), but the microbial traits that facilitate this process are unresolved. While current theory and biogeochemical models suggest microbial carbon-use efficiency and growth rate are positive predictors of SOC, recent observations demonstrate these relationships can be positive, negative, or neutral. To parse these contradictory effects, we used a ^13^C-labeling experiment to test whether different microbial traits influenced the transformation of plant C into SOC within the microbial habitats surrounding living root inputs (rhizosphere) versus decaying root litter (detritusphere), under both normal soil moisture and droughted conditions. In the rhizosphere, bacterial-dominated communities with fast growth, high carbon-use efficiency, and high production of extracellular polymeric substances formed microbial-derived SOC under normal moisture conditions. However, in the detritusphere – and the rhizosphere under drought – more fungal-dominated communities with slower growth but higher exoenzyme activity formed plant-derived SOC. These findings emphasize that microbial traits linked with SOC accrual are not universal, but contingent on how microorganisms allocate carbon under different resource conditions and environmental stressors.

## MAIN

Soil microorganisms shape the global carbon balance by transforming plant C inputs to soil organic carbon (SOC) – the largest actively-cycling pool of C in the terrestrial biosphere (>1500 Pg C)^1^. It is crucial to understand which soil microbial traits influence SOC accrual under climate change^2^ – particularly mineral-associated SOC, the largest and slowest-cycling C pool in Earth’s mineral soils, since its fate will substantially influence atmospheric CO_2_ concentrations and the trajectory of planetary warming^3^. Decaying microbial residues – such as cell envelopes, DNA, and extracellular polymeric substances (collectively described as ‘microbial necromass’) – are increasingly recognized as key ingredients of mineral-associated SOC, and can comprise 50% or more of the total pool^4–6^. A dominant hypothesis is therefore that greater microbial carbon-use efficiency (CUE) and faster growth and turnover should lead to greater microbial necromass production, and greater accrual of mineral-associated SOC^7,8^. While this hypothesis has gained traction – and is represented in several ‘microbial-explicit’ biogeochemical models^9,10^ – few empirical studies have directly tested it. Those that have yielded mixed support. In ecosystems like croplands and grasslands, where a greater proportion of SOC appears to be ‘microbial-derived’ (i.e., plant C that is microbially assimilated and transformed to various microbial compounds, which then associate with soil minerals), greater microbial growth and CUE can lead to more mineral-associated SOC^11^. However, in ecosystems like temperate forests where a greater proportion of SOC appears to be ‘plant-derived’ (i.e., plant C that may be partially decomposed by microbial exoenzymes into simpler plant compounds, but does not pass through a microbial body before associating with soil minerals), these relationships can be neutral or negative^7^.

Growth rate and CUE therefore have contradictory effects on mineral-associated SOC formation, in part due to whether mineral-associated SOC is plant-derived or microbial-derived. However, the mechanisms underpinning these patterns remain elusive, impeding empirical understanding of how microbes control SOC cycling, and the subsequent development of accurate SOC models that predict SOC persistence and its responsiveness to climate change^2,7,12^. To understand contrasting effects of microbial traits on mineral-associated SOC, it is necessary to untangle their effects at the scale of the microbial habitat – the site where different sources of plant C input are transformed to SOC^12^. The two primary and contrasting sources of plant C input to the mineral soil are: (i) lower-molecular weight living root inputs (rhizodeposits like root exudates), which support a microbial habitat in the zone surrounding living roots known as the ‘rhizosphere’, and (ii) more biochemically complex decaying root litter, which creates a microbial habitat known as the ‘detritusphere’^13–15^. Differences in the biochemical composition, quantity, and rate of delivery of rhizodeposits versus root litter select for distinct microbial communities that specialize on these different substrates^16,17^. By extension, these distinct microbial communities should express different sets of traits that facilitate processing of these C inputs, potentially leading to different relationships between these traits and mineral-associated SOC accrual and composition in the rhizosphere and detritusphere^12^.

In the rhizosphere, fast-growing microorganisms with high CUE should thrive, due to the abundance of lower molecular weight, easily-assimilated substrates that can be biosynthesized into intracellular microbial biomass and extracellular polymeric substances^18^. As these microbial compounds become the ingredients in microbial-derived, mineral-associated SOC, there should be a positive relationship in the rhizosphere between mineral-associated SOC and traits responsible for generating microbial necromass, like growth rate, CUE, and high biomass yield^12^. In the detritusphere, however, microorganisms with fast growth and high CUE may not be as competitive on more complex sources of organic C (e.g., root detritus), where greater exoenzyme activity is required to decompose these complex compounds into simpler plant compounds^19^. Since exoenzyme activity often trades off with fast growth and high yield^18,20–22^, slower growth but greater exoenzyme activity may be better predictors of mineral-associated SOC in the detritusphere. Mineral-associated SOC formed from root detritus may also be more plant-derived, if a substantial portion of these partially decomposed, plant compounds directly associate with minerals without undergoing microbial assimilation^23,24^. These posited trait patterns in the rhizosphere and detritusphere may be also modulated by key soil variables that can affect the expression of traits related to mineral-associated SOC. Altered soil moisture, for example, is one dominant effect of climate change on soil microbial community composition and function^25^, and can impact the expression of traits like CUE and exoenyzme activity^26^.

Here, we conducted a ^13^C-labeling experiment to test whether divergent microbial traits and taxa influenced mineral-associated SOC accrual and chemistry in the rhizosphere and detritusphere, and determine how soil moisture modulated these relationships. At three points during a 12-week spring growing season of the annual grass *Avena barbata*, we tracked the movement of ^13^C-labeled rhizodeposits and ^13^C-labeled root litter into microbial communities and mineral-associated SOC of the rhizosphere and detritusphere, respectively, under both normal moisture (~16% gravimetric soil moisture)^27^and droughted soil conditions (~8% soil moisture) of a semi-arid grassland soil^28,29^. We measured (i) the accrual and chemical composition of mineral-associated SOC, (ii) a suite of microbial traits (community-level carbon-use efficiency, growth rate, microbial biomass, extracellular polymeric substances, extracellular enzyme activity), and (iii) taxon-specific microbial activity via ^13^C quantitative stable isotope probing. We hypothesized that distinct microbial traits and taxa in the rhizosphere and detritusphere would be correlated with ^13^C-mineral-associated SOC accrual and chemical composition (i.e., plant-derived versus microbial-derived), and these relationships would be shaped by soil moisture. Such evidence for divergent traits in the rhizosphere versus detritusphere would both help clarify contradictory observations surrounding the importance of growth rate and CUE in mineral-associated SOC cycling, and emphasize that future theoretical and modeling efforts should include how microbial trait trade-offs shape which traits are linked with SOC accrual in different soil contexts^12,18,30^.

### Microbial traits and mineral-associated SOC in the rhizosphere and detritusphere

To identify relationships between microbial traits and mineral-associated SOC formed in the rhizosphere and detritusphere (Figs. 1–2), we measured ^13^C mineral-associated SOC derived from ^13^C-rhizodeposits versus ^13^C-root litter, as well as several key microbial traits at 4, 8, and 12 weeks of a single growing season of *A. barbata* (Supp. Fig. S1-S2). These traits included microbial community-level growth rate and carbon use efficiency (CUE), which influence the amount of microbial necromass produced; total microbial biomass C and extracellular polymeric substances (EPS), which reflect dominant pools of microbial C that can form microbial-derived SOC^12,31,32^; and cumulative exoenzyme activity, which influences the extent to which a microbial community can depolymerize complex substrates (for example, root litter) into simpler compounds^33^. We focused primarily on week 12 measurements, as this time point had the most pronounced drought treatment effects (Supp. Fig. S3), as well as maximum accrual of ^13^C-mineral-associated SOC (Supp. Table S1).

Overall, we found that the detritusphere selected for a distinct set of microbial traits relative to the rhizosphere, this pattern was modulated by soil moisture (Fig. 1; Supp. Table S2). By week 12, the detritusphere microbial community had ~35% greater cumulative exoenzyme activity relative to the rhizosphere community, averaged across both moisture treatments (Fig. 1a; Tukey HSD, p < 0.05). In contrast, the rhizosphere microbial community had a faster community-level growth rate (Fig. 1b), higher CUE (Fig. 1c) and greater microbial biomass C (Fig. 1d) than the detritusphere community under normal moisture (Tukey HSD; p < 0.05). Under droughted conditions, however, the growth rate, CUE, and microbial biomass C in the rhizosphere were significantly reduced, and not statistically different from the detritusphere (Fig. 1b-d; Tukey HSD, p > 0.05).

**Figure 1.**
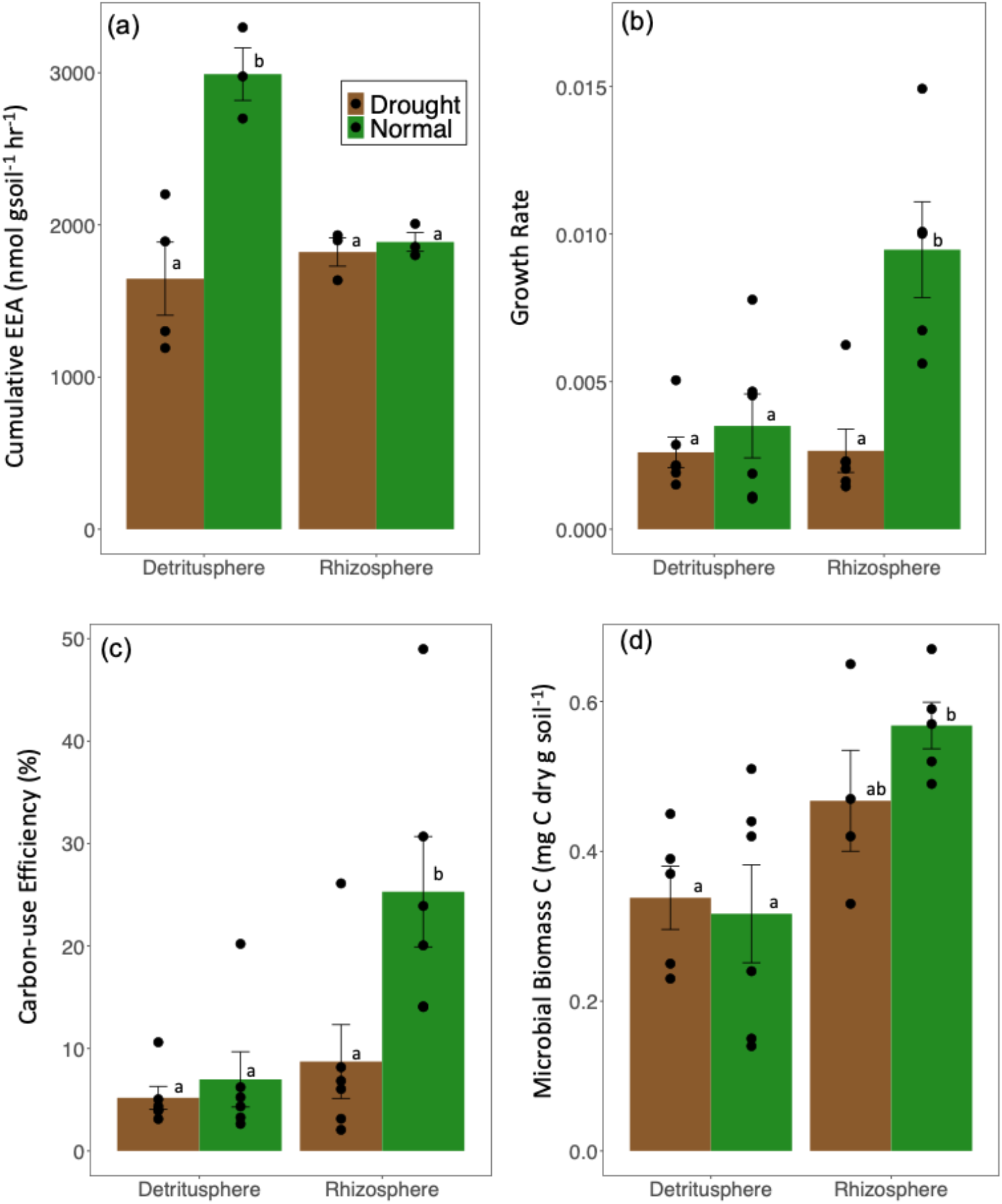
Microbial traits in the rhizosphere and detritusphere of *Avena barbata* after 12 weeks of growth/incubation in a California annual grassland soil under normal moisture (green) or droughted conditions (brown). Traits include: (a) cumulative exoenzyme activity (EEA; *n* = 4), (b) microbial community-level growth rate (*n* = 6), (c) microbial community-level carbon use efficiency (CUE; *n* = 6), and (d) microbial biomass carbon (MBC; *n* = 6). Letters indicate significant differences between means determined via Tukey’s HSD (p < 0.05). For full model output for regressions including all three time points (weeks 4, 8, 12) see Supp. Table S3.

We also observed rhizosphere vs. detritusphere contrasts in the relationships between microbial community traits and ^13^C-mineral-associated SOC (Fig. 2). In the rhizosphere, across all time points, community-level growth rate and ^13^C-EPS were both significantly positive predictors of ^13^C-mineral-associated SOC (p < 0.001; Supp. Table S3). By week 12, growth rate, CUE, ^13^C-EPS, and ^13^C-microbial biomass C were all positively correlated with ^13^C-mineral-associated SOC in the rhizosphere (Fig. 2a; p < 0.02). However, in the detritusphere, none of these four traits were significant predictors of mineral-associated SOC, whether analyzed across all time points or solely at week 12 (Fig. 2b, Supp. Table S4, p > 0.5).

**Figure 2.**
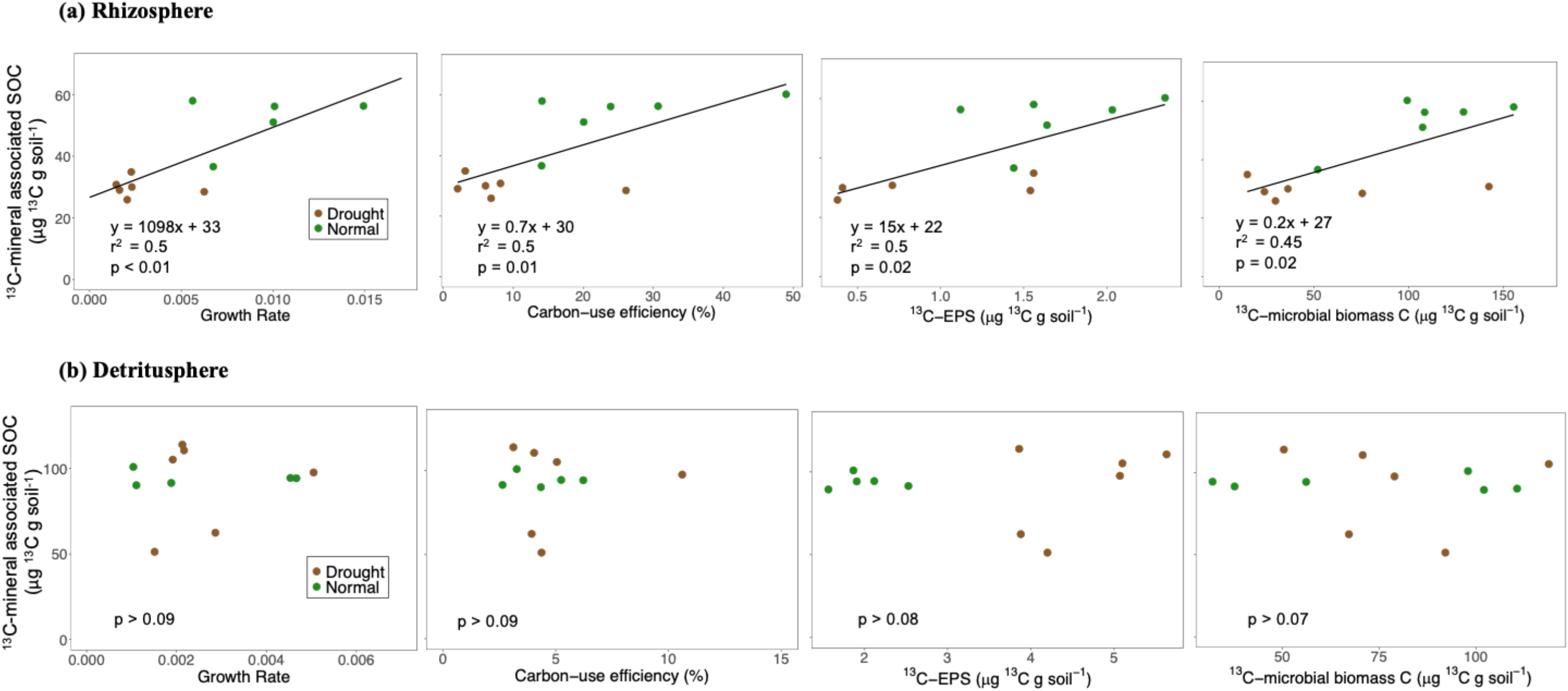
Linear regressions between different microbial traits and ^13^C-mineral-associated soil organic carbon (SOC; ug ^13^C g soil ^−1^) formed from: (a) ^13^C-rhizodeposits in the rhizosphere and (b) ^13^C-root detritus in the detritusphere in an annual grassland soil. Microcosms were maintained under either normal moisture (green) or droughted conditions (brown) for 12 weeks. Traits include, from left to right, community-level growth rate, carbon-use efficiency (CUE), ^13^C-extracellular polymeric substances (EPS), and ^13^C-microbial biomass C (*n* = 12). Notably, the scale of the y-axes are different between rhizosphere and detritusphere data plots due to different amounts of ^13^C-rhizodeposits versus ^13^C-root litter entering the soil.

### Chemical composition of mineral-associated SOC

To capture differences in the chemical composition of mineral-associated SOC formed in the rhizosphere and detritusphere, we used both: (i) ^13^C-nuclear magnetic resonance (^13^C-NMR), which provides chemical composition of mineral-associated at the bulk scale, and (ii) scanning transmission X-ray microscopy in combination with near edge X-ray absorption fine structure spectroscopy (STXM-NEXAFS), which yields spatial chemical composition data of SOC at the microscale.

^13^C-NMR spectra of ^13^C-mineral-associated SOC indicated a more microbial-derived signature in the rhizosphere relative to the detritusphere, as measured via the ratio of alkyl/O-alkyl regions (A/OA ratio; 0-45/45-110 ppm) (Fig 3a-b; Supp. Table S5). The A/OA ratio reflects the degree of microbial decomposition of organic carbon: a higher relative ratio indicates greater microbial transformation of ^13^C plant C, whereas a lower ratio points to more intact ^13^C plant C^34,35^. The A/OA ratio was 32% higher in the rhizosphere than the detritusphere under normal moisture conditions (Fig. 3a), and 16% higher under droughted conditions (Fig. 3b).

**Figure 3.**
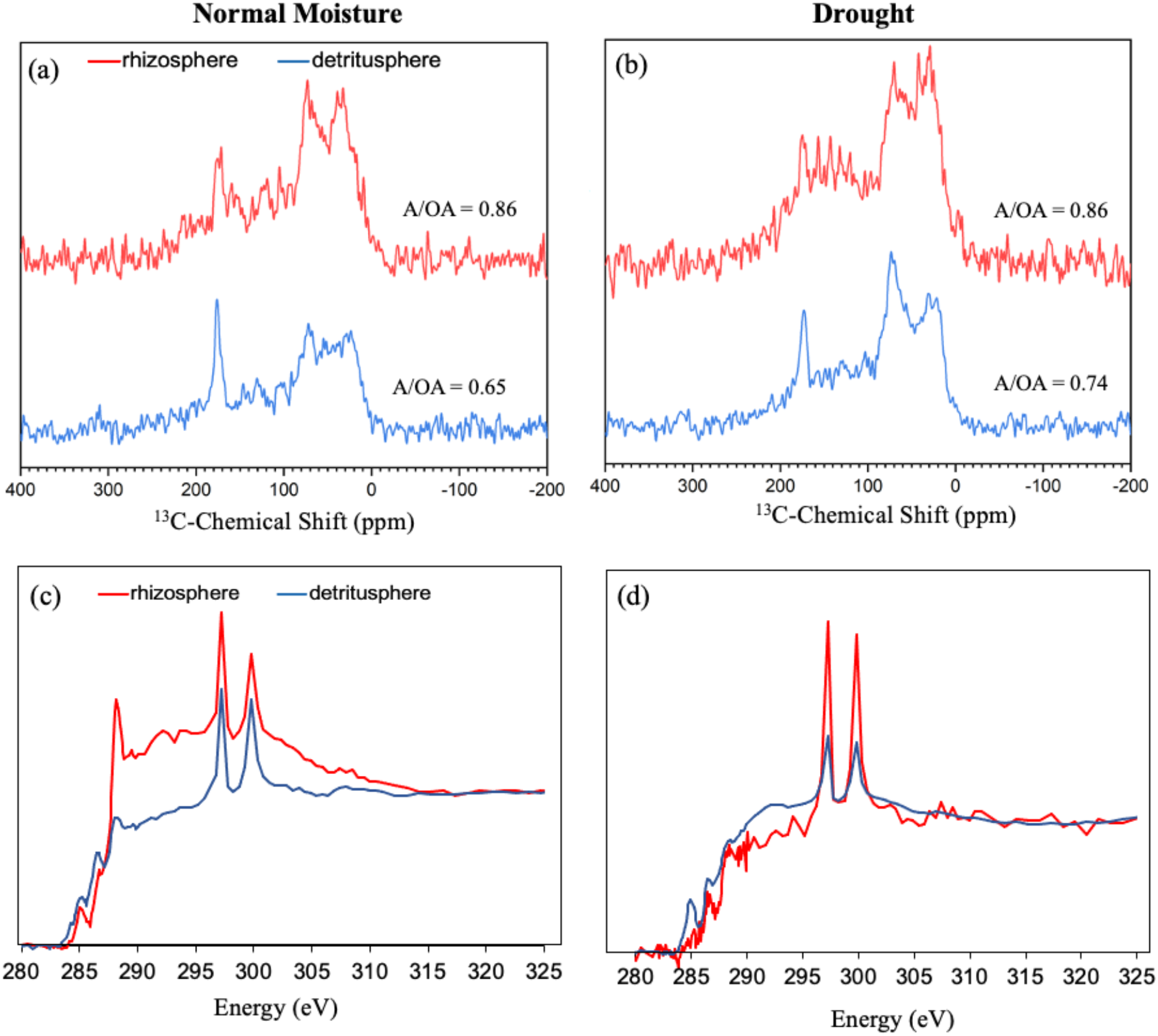
Chemical composition of mineral-associated soil organic carbon (SOC) in the rhizosphere (red) and detritusphere (blue) at week 12, measured via: (a,b) ^13^C-nuclear magnetic resonance (^13^C-NMR), and (c,d) scanning transmission X-ray microscopy in combination with near edge X-ray absorption fine structure spectroscopy (STXM-NEXAFS). A/OA = the ratio of alkyl C to O-alkyl C (0-45/45-110 ppm); a higher ratio indicates greater microbial transformation of organic matter^35^. STXM-NEXAFS spectra are averages of 3-5 spectra performed on a single replicate, ^13^C-NMR was conducted on a composite sample of 6 replicates (n = 6) per treatment.

In the normal moisture soils, the C NEXAFS spectra of rhizosphere mineral-associated SOC had a distinctive peak around 288.5 eV in the carboxyl region and smaller peaks in the 284.7 and 286.7 eV positions (Fig. 3c). Overall, we observed a more oxidized profile in the rhizosphere mineral-associated SOC under normal moisture, consistent with more ‘microbial-derived’ signature and reflecting microbial assimilation and transformation of plant C compounds. In contrast, detritusphere mineral-associated SOC had a less oxidized, more ‘plant-derived’ signature, indicating a lower proportion of microbially-assimilated detrital C. Under drought conditions, however, the spectra of mineral-associated SOC formed in the rhizosphere was more plant-derived – and appeared similar to mineral-associated SOC formed in the detritusphere (Fig 3d). Put together, our ^13^C-NMR and STXM-NEXAFS data indicate that at both a bulk scale and microscale, there was a greater proportion of microbial-derived, mineral-associated SOC in the rhizosphere, and a greater proportion of more plant-derived, mineral-associated SOC in the detritusphere^23^; and this pattern was modulated by drought.

### Microbial activity in the rhizosphere and detritusphere

To assess the roles of different microbial taxa on the formation of mineral-associated SOC, we measured bacterial and fungal relative abundance in the rhizosphere and detritusphere by quantifying total 16S and 18S rRNA gene copies at week 12 via quantitative PCR. Bacteria were ~41% more abundant in the rhizosphere relative to the detritusphere, whereas fungi were ~65% more abundant in the detritusphere than the rhizosphere (Supp. Fig. S4). We used ^13^C quantitative stable isotope probing^36^ (^13^C-qSIP) to determine whether these differences translated to distinct patterns of active ^13^C-assimilation amongst bacterial and fungal taxa. We measured both the cumulative bacterial and fungal assimilation of ^13^C in the rhizosphere and detritusphere (i.e., the sum total of each taxon’s atom fraction excess ^13^C value weighted by its relative abundance), as well as the proportional ^13^C assimilation by different fungal and bacterial amplicon sequence variants (ASVs) (a standardized metric of ^13^C assimilation by individual taxa^37^; see Methods).

Cumulative bacterial and fungal ^13^C assimilation measurements revealed that in the detritusphere, fungal ^13^C assimilation was 80% greater than bacterial ^13^C assimilation under normal moisture conditions, and 46% greater under drought (Fig. 4b; p < 0.05). In the rhizosphere, bacterial ^13^C assimilation was 28% greater than fungal assimilation under normal moisture conditions (Fig. 4a; p < 0.1). Under droughted conditions, however, bacterial and fungal ^13^C assimilation in the rhizosphere were roughly equivalent (Fig. 4a; p > 0.1). Together, this suggests that in the detritusphere, greater fungal activity primarily explained the trait patterns we observed, including greater exoenzyme activity and slower growth; whereas in the rhizosphere, higher bacterial activity explained more of the observed trait patterns of faster growth, higher CUE, and greater biomass (Fig 1). In the rhizosphere, drought modulated this trend (Fig 4a-b) – echoing the observations noted above that trait expression and the chemical composition of mineral-associated SOC in the rhizosphere are tempered under drought (Fig 1; Fig. 3b).

**Figure 4.**
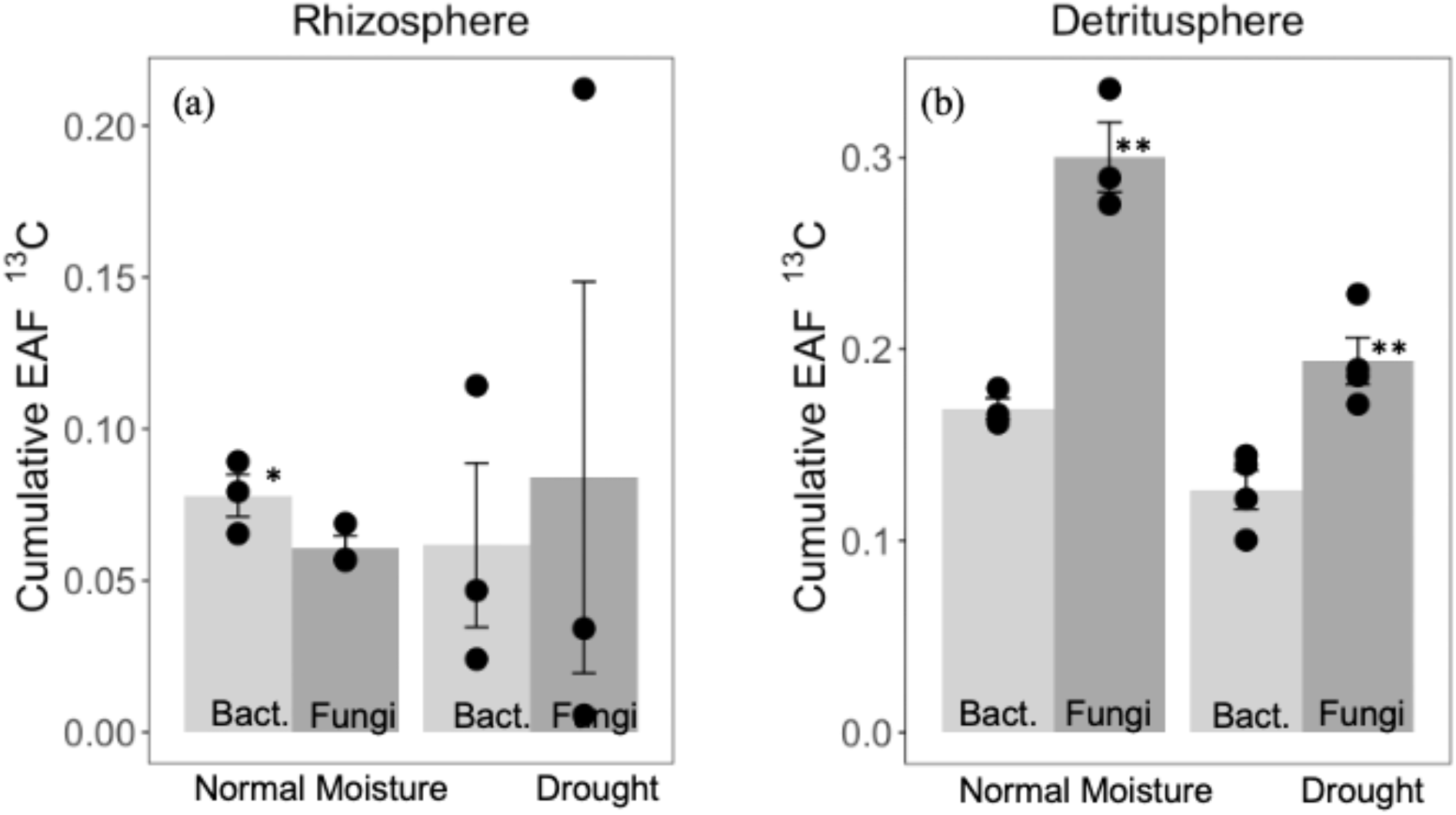
Cumulative ^13^C-assimilation of bacteria (‘bact.’) and fungi after 12 weeks in the (a) rhizosphere and (b) detritusphere of an annual grassland soil, measured via ^13^C-quantitative stable isotope probing. Cumulative excess atom fraction (EAF) ^13^C represents the summed EAF ^13^C value for each taxa, weighted by its relative abundance. Asterisks indicate significant differences between means of bacterial and fungal cumulative EAF ^13^C in each soil habitat × moisture treatment combination, measured via t-tests (* p < 0.1, ** p < 0.05). *N* = 4. Error bars indicate standard error.

In both the rhizosphere and detritusphere, a relatively small number of bacterial or fungal ASVs accounted for the majority of ^13^C assimilation (Fig. 5). Within each microbial habitat × moisture treatment, between 7–13 bacterial ASVs and 3–6 fungal ASVs accounted more than 50% of proportional ^13^C-assimilation (hereon we define these as ‘top assimilators’; see Methods) (Supp. Table S6). These top bacterial assimilators represented only 3–4% of all bacterial ASVs that assimilated ^13^C, and top fungal assimilators represented only 4–5% of all fungal ASVs that assimilated ^13^C (Supp. Table S6). Given the outsized role played by a small number of microbial taxa in assimilating ^13^C, we queried the functional traits of these top assimilators (grouped by family) through literature searches to explore the relationships between microbial traits and mineral-associated SOC formation.

**Figure 5.**
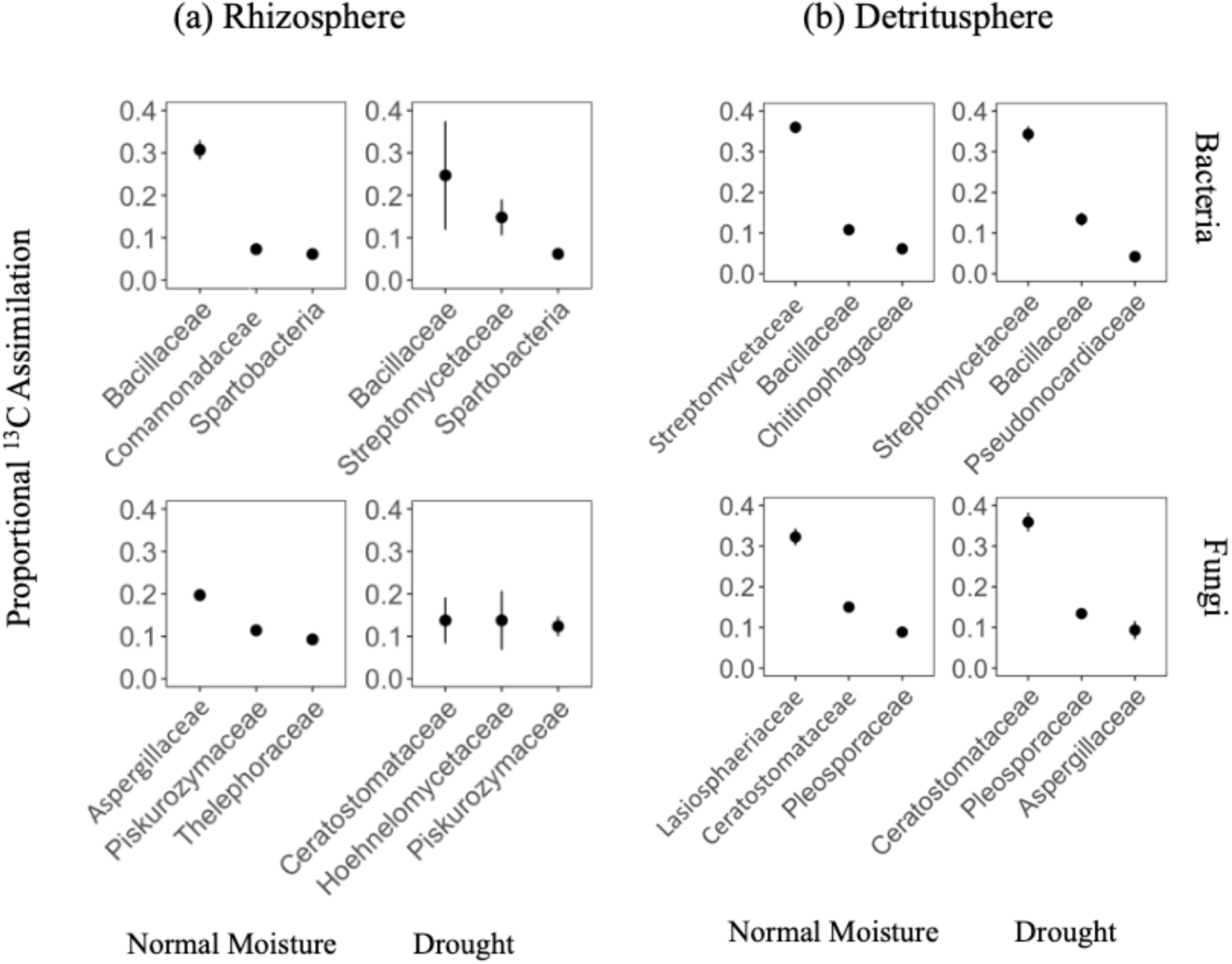
Taxon-specific proportional ^13^C assimilation of top fungal and bacterial ^13^C assimilating Amplicon Sequence Variants (ASVs) found in the (a) rhizosphere and (d) detritusphere (*n* = 4) from an annual grassland soil incubated under normal moisture and droughted conditions. As discussed in Methods, ‘top assimilators’ are defined as microbial taxa that together account for > 50% of proportional ^13^C assimilation (see Supp. Table S6). ASVs were grouped by family; the three families with the highest ^13^C proportional assimilation values are shown below; additional top assimilators are shown in Supp. Table S6. *N* = 4. Error bars indicate standard error.

In the detritusphere, ^13^C assimilation was dominated by filamentous, saprotrophic microorganisms with high lignocellulolytic potential (Fig. 5b). The top fungal assimilators include families in the Ascomycota with filamentous growth forms (such as *Ceratostomataceae, Lasiosphaeriaceae*, and *Pleosporaceae)*, all of which are capable of producing extracellular enzymes to degrade complex C compounds including lignin^38^. Compared to these fungal taxa, bacteria contributed quantitatively less to ^13^C assimilation in the detritusphere (Fig. 5b); top bacterial assimilators also included lignocellulolytic taxa, with both filamentous (*Streptomycetaceae, Bacillaceae*) and non-filamentous (*Chitinophagaceae*) growth forms.

In the rhizosphere, top bacterial assimilators in normal moisture soils were from families in the Proteobacteria (*Comamonadaceae, Bradyrhizobiaceae*), Firmicutes (*Bacillaceae*), and Verrucomicrobia (*Spartobacteria*) (Fig. 5a). Broadly, these bacterial families are known to be successful rhizosphere competitors, exhibiting traits like high rates of motility, secondary metabolite production, and production of antifungal compounds^39–42^. The top rhizosphere fungal assimilators were mostly saprotrophs, from the families *Aspergillaceae*, *Theloporaceae*, and *Piskurozymaceae*.

Droughted conditions significantly affected which taxa were responsible for ^13^C assimilation in the rhizosphere (Supp. Fig. S5; Permanova, p < 0.05). Notably, there was significant overlap in the taxa that were top rhizosphere ^13^C assimilators under drought, and taxa that were top assimilators in the detritusphere (regardless of soil moisture). In droughted rhizosphere soils, a filamentous *Streptomycetaceae* from the Actinobacteria became a top assimilator (Fig 5a, Supp. Table S6). *Streptomycetaceae* grow substrate mycelium – a growth form that is posited to be advantageous for withstanding moisture stress and colonizing detritus^43,44^. *Streptomycetaceae* were also dominant assimilators in the detritusphere, possibly due to their capacity to produce non-catalytic binding proteins and hydrolytic enzymes to degrade complex organic matter, and various antibiotics that may help them compete with soil fungi^45^. Top fungal assimilators in the moisture-limited rhizosphere were also dominant in the detritusphere, including *Aspergillaceae*, *Ceratostomataceae*, and *Piskurozymaceae* (Fig. 5b).

## DISCUSSION

Increasing evidence suggests that soil microorganisms play a central role in the formation of mineral-associated SOC, and empirical and modeling efforts are keen to identify which microbial traits are linked with SOC accrual, especially under a changing climate^2,5,9,26,46^. Climate change factors like altered precipitation, warming, and elevated CO_2_ can alter the amount and chemical composition of plant C inputs like root exudates, the composition and dominant traits of soil microbial communities, as well as the persistence and chemistry of mineral-associated SOC^25,26^. Much of the effort to link microbial traits with SOC accrual has focused on community-level CUE and growth rate, because these traits are posited to be important for the formation of microbial-derived, mineral-associated SOC that is formed from the intracellular and extracellular residues of microorganisms that associate with soil minerals after cell death and turnover^5,8,9,11^. However, relatively few controlled studies have tested the role of these traits in different soil habitats or environmental contexts, or accounted for other pathways of SOC formation that may predominate. For example, the partial decomposition of complex plant compounds by microbial exoenzymes into simpler plant compounds – which associate with minerals to form plant-derived SOC – should select for a different set of microbial traits. Indeed, in our study, we found that that two dominant and contrasting types of plant C input to the mineral soil – rhizodeposits and root litter – selected for divergent sets of microbial traits which mapped onto contrasting microbial life history strategies and pathways of mineral-associated SOC formation in the rhizosphere and detritusphere.

In the rhizosphere, the abundance of lower-molecular weight, easily-assimilated C substrates, selected for microorganisms with faster and more prolific growth (Fig. 1). Overall, a fast-growing, efficient, and high biomass and high EPS-producing rhizosphere microbial community transformed rhizodeposits into mineral-associated SOC under normal moisture conditions (Fig. 2a). Together, these traits typify a microbial life history strategy specialized for environments with high availability of simple resources, like sugars and amino acids (described recently as a ‘high yield’ strategy^18^). ^13^C-qSIP indicated that bacteria assimilated more ^13^C than fungi in the rhizosphere under normal moisture conditions (Fig 4a), and taxa known to be rhizosphere-dwelling in the *Bacillaceae*, *Bradyrhizobiaceae*, and *Comamondaceae* were particularly active (Fig. 5a). On average, soil bacteria are recognized to exhibit growth and turnover rates that are an order of magnitude faster than soil fungi^47,48^. This effect that may be even more pronounced for the competitive rhizosphere bacteria we measured, since the rhizosphere is a zone of high predation and turnover, and thus a large yield of microbial necromass can be generated^49,50^. The chemical composition of mineral-associated SOC in the rhizosphere under normal moisture had a more microbial-derived signature than the detritusphere, as determined by STXM-NEXAFS and ^13^C-NMR (Fig. 3). This suggests that a greater proportion of mineral-associated SOC in the rhizosphere was formed via microbial assimilation and transformation of simple plant C substrates into microbial biomass and residues, which then became associated with soil minerals (described previously as the ‘*in vivo* microbial turnover pathway’^51^).

In the detritusphere, the dominant microbial traits and active taxa were quite different from those we had identified in the rhizosphere. The more complex, polymeric, detrital compounds selected for a more fungal-dominated community, with slower growth, lower biomass and CUE, but greater extracellular enzyme activity. This set of microbial traits, which were recently described as part of a ‘resource acquisition’ life history strategy^18^, are thought to trade-off with fast and efficient growth^20^. ^13^C-qSIP revealed that microbial decomposition in the detritusphere was dominated by filamentous fungi (*Ceratostomataceae, Lasiosphaeriaceae*, and *Pleosporaceae*), which are known to not only have an average turnover time that is much longer than soil bacteria (as discussed above), but can also exude high quantities of extracellular enzymes to degrade complex organic matter in litter or SOC^52^ (Fig. 5). Indeed, recent work has shown that extracellular enzyme production and growth rate trade-off in filamentous fungi^53^. The chemical composition of mineral-associated SOC in the detritusphere had a more plant-derived signature than the rhizosphere, suggesting that a greater proportion of SOC was formed via partial decomposition of complex plant compounds into simpler plant compounds that directly associated with soil minerals (described previously as the ‘*ex vivo* modification pathway’^51^).

Soil moisture played a key moderating role in the rhizosphere. Drought selected for a microbial community in the rhizosphere that more closely resembled the detritusphere community – both in the traits expressed, as well as the identity of active taxa assimilating ^13^C at week 12. Under drought conditions, the growth rate, CUE, and microbial biomass of the rhizosphere community were reduced to levels comparable with the detritusphere (Fig. 1). ^13^C-qSIP indicated that active taxa in the droughted rhizosphere at week 12 were similar to the detritusphere, particularly filamentous fungi and filamentous bacteria (e.g. *Streptomycetes*) with lignocellulytic capabilities. This bolsters prior long-standing hypotheses and some recent work^54–56^ showing that these taxa may be more resilient to drought conditions. In addition, STXM-NEXAFS spectra indicated a more plant-derived signature of mineral-associated SOC in the rhizosphere under drought, comparable with the detritusphere (Fig. 3c). This may have been due to greater root death and turnover under drought, or drought-induced shifts in the chemical composition of rhizodeposits (i.e., greater amounts of complex organic acids and other compounds)^57^, which can affect which taxa are active and the traits they express^58^.

Put together, several lines of converging evidence (microbial traits, ^13^C-qSIP, chemical composition of SOC) suggest that: (1) distinct pathways of mineral-associated SOC formation predominate in the rhizosphere versus the detritusphere, (2) these pathways are associated with distinct microbial traits and taxa, and (3) soil moisture modulates these relationships (Fig. 6). The first formation pathway (‘*in vivo* microbial turnover’; Fig. 6a) was dominant in the rhizosphere, characterized by microbial assimilation and biosynthesis of lower-molecular weight plant inputs (i.e., root exudates and other rhizodeposits) into different intracellular and extracellular microbial compounds, which subsequently formed microbial-derived, mineral-associated SOC (Fig. 6a). Microbial traits like growth rate, CUE, and EPS were therefore positive predictors of mineral-associated SOC accrual in the rhizosphere. The second pathway (‘*ex vivo* modification’; Fig. 6b) likely dominated in the detritusphere, and is characterized by exoenzyme-mediated depolymerization of more complex inputs (i.e., root litter) into simpler plant compounds, which formed plant-derived, mineral-associated SOC. In the detritusphere, CUE, growth rate, and microbial biomass were not significantly associated with mineral-associated SOC (Fig. 2); instead, root litter selected for traits like greater extracellular enzyme activity and slower growth. Drought modulated these trends, by selecting for a rhizosphere microbial community that behaved more like the detritusphere community in terms of which taxa were active in assimilating ^13^C, the community-level traits expressed, as well as the formation of mineral-associated SOC via *ex vivo* modification.

**Figure 6.**
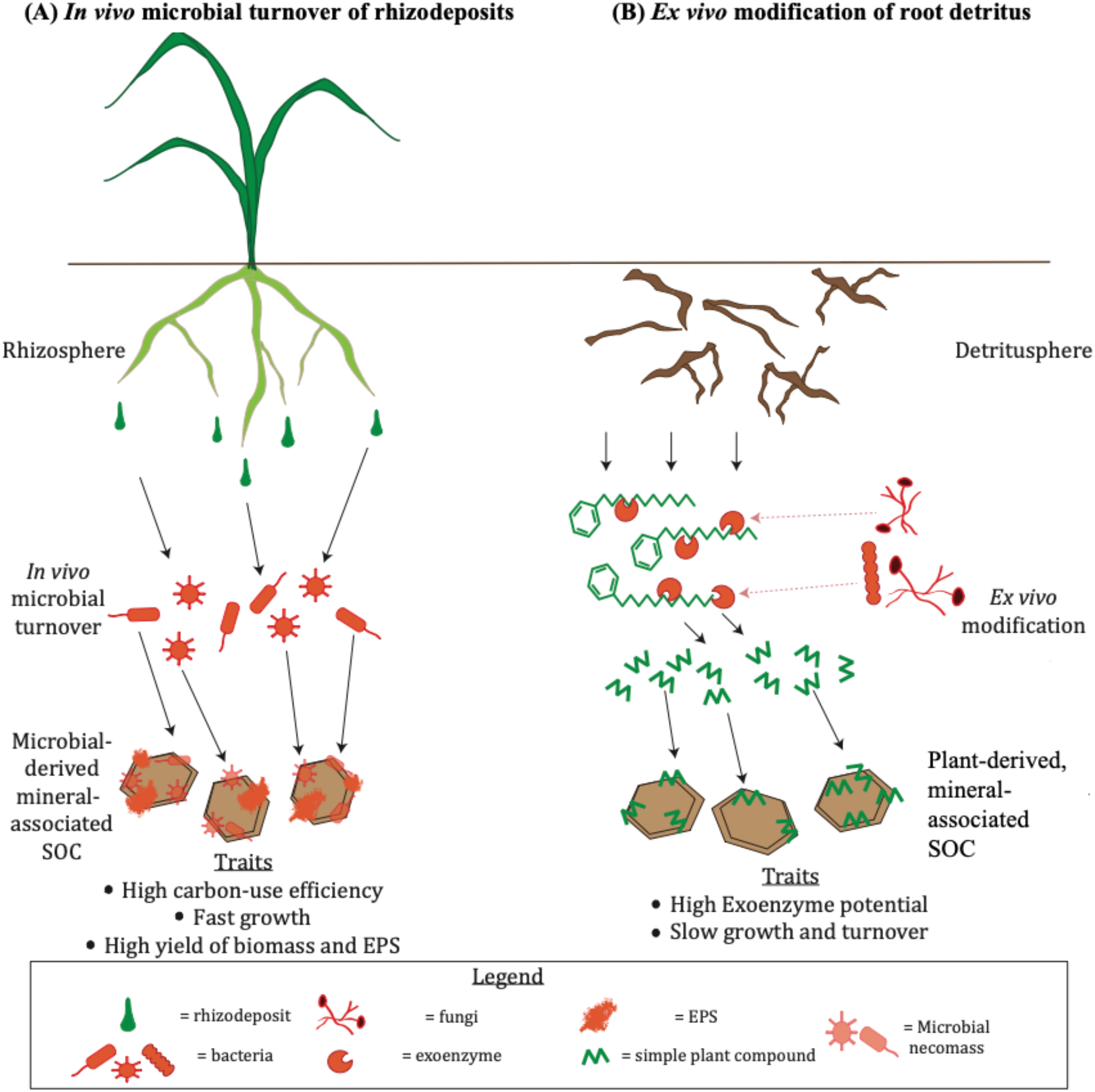
Conceptual model illustrating divergent microbial traits and taxa associated with distinct pathways of mineral-associated SOC formation from rhizodeposits versus root detritus of the annual grass *Avena barbata*. (A) In the rhizosphere, lower-molecular weight rhizodeposits were microbially assimilated and transformed (the ‘*in vivo* microbial turnover pathway’) by a more active bacterial community to form microbial-derived, mineral-associated SOC. (B) In the detritusphere, complex root detrital compounds were partially decomposed into simpler plant compounds by microbial exoenzymes (the ‘*ex vivo* modification pathway) by a more active fungal community to form plant-derived, mineral-associated SOC. Drought modulated this trend in the rhizosphere, selecting for a microbial community and pathway of SOC formation that more closely resembled the detritusphere.

It is critical to understand which microbial taxa and traits are linked with different pathways of mineral-associated SOC formation, since these pathways can lead to SOC pools with different chemical compositions, and degrees of persistence and responsiveness to climate change^26^. Our results challenge current hypotheses of a consistent, positive relationship between mineral-associated SOC and traits like CUE, growth rate, and microbial biomass^8^. While we observed positive relationships between these traits and mineral-associated SOC in the rhizosphere, the lack of significant relationships in the detritusphere demonstrate that CUE and growth rate may be uncoupled from mineral-associated SOC formation in certain contexts, likely due to allocation of C to other compounds, such as extracellular enzymes^7,59^. These findings can help explain prior contrasting observations on the role of CUE and growth rate in mineral-associated SOC accrual^7,11^. Our results also emphasize that it is important to consider how soil microorganisms allocate their C under different resource environments and different climate stressors – whether to growth, resource acquisition, or to products like storage or stress compounds – in order to understand which microbial traits and taxa will be associated with the accrual of mineral-associated SOC^2,9,10^. Incorporating such a context-specific understanding into theory and microbial-explicit biogeochemical models will be necessary to accurately depict how microorganisms influence the global carbon balance under climate change.

## METHODS

### Experimental Design

We conducted a ^13^C-labeling greenhouse tracer and soil moisture manipulation study, where we maintained planted and unplanted microcosms inside growth chambers for 12 weeks, to track *Avena barbata* rhizodeposits versus root detritus, respectively (Supp. Fig. S1-S2). Soil for the experiment was collected from 0-10 cm depth (‘A’ mineral horizon) below a stand of *A. barbata* at the University of California Hopland Research and Extension Center (39°00.106`N, 123°04.184`W), which is the traditional and ancestral territory of the Shóqowa and Hopland People. The site is a Mediterranean annual grassland ecosystem (MAT max/min = 23/7 C; MAP = 956 mm yr^−1^), where *Avena* spp. is the dominant vegetation. The soil is classified as a Typic Haploxeralfs of the Witherall-Squawrock complex, soil pH is ~5.6, contains 45% sand, 36% silt, and 19% clay^27^. Initial C and N content were 2.2% and 0.24%, respectively.

After collection, roots were removed, and soil was passed through a 2-mm sieve and packed to field bulk density (1.21 g cm^3^) in rectangular acrylic microcosms (11.5 × 2.9 × 25.5 linear cm), as described previously^60^. For planted microcosms (‘rhizodeposit’ treatment), three germinated *Avena barabata* seedlings (collected from Hopland in spring 2018) were planted evenly apart in the soil, to simulate field seedling density. Microcosms were then incubated inside 8 growth chambers (56 × 56 × 76 linear cm) with a 16-h light period per day (from 6 a.m. to 10 p.m.) and a maximum daytime and nighttime air temperature of 27°C and 24°C, respectively. A subset of planted microcosms were grown inside ^13^CO_2_ growth chambers (filled with ~99 atom% ^13^CO_2_, *n* = 6 per time point × moisture treatment combination; Supp. Fig. S1), in order to track ^13^C-labeled rhizodeposits into the rhizosphere. A control set of planted microcosms (*n* = 4) grown in ^12^CO_2_ growth chambers (with a natural abundance atmosphere) was used for quantitative stable isotope probing (discussed below) and to increase available soil for measurements that did not require a ^13^C-label (for example, exoenzyme activity). ^12^CO_2_ chambers were monitored using an Infrared Gas Analyzer (IRGA; SBA-5 CO_2_ Gas Analyzer, PP systems, Amesbury, MA, USA) and ^13^CO_2_ chambers were monitored using a Picarro G2201-I cavity ringdown spectrophotometer (Picarro Santa Clara, CA, USA). The entire system was controlled with a CR1000 datalogger (Campbell Scientific Logan, UT, USA). ^12^CO_2_ and ^13^CO_2_ chambers were monitored in parallel to each other; each chamber was scanned once every 40 minutes to monitor CO_2_ concentrations, and customized code (CRBasic) was developed to calculate the amount of CO_2_ to add to each chamber based on CO_2_ drawdown and photosynthetically active radiation (PAR).

In unplanted microcosms (‘root detritus’ treatment), a 28-μm mesh ‘detritusphere’ bag containing ~65 g of Hopland soil mixed with ~1-5 mm fragments of *A. barbata* root detritus (0.013 g detritus dry g soil^−1^) was buried in the microcosm center (Supp. Fig S2)^16^. An ^13^C-enriched subset (*n* = 6) contained ^13^C-labled root detritus (77 ± 1.7 atom% ^13^C-labeled), while a control subset (*n* = 4) contained natural abundance root detritus (1.1 atom% ^13^C). Unplanted microcosms were incubated within growth chambers in a natural abundance CO_2_ environment for the experimental period.

Planted and unplanted microcosms were maintained at one of two moisture treatments, to simulate differences in soil moisture during the spring growing season in California semiarid grasslands: ‘normal moisture’ (~16% ± 0.3 gravimetric soil moisture; mean ± standard error) or ‘spring drought’ conditions (~8% ± 0.5 gravimetric soil moisture)^27^. Soil moisture was monitored throughout the experimental period by weighing all microcosms twice weekly, and adjusting soil moisture by mass^60^. At each harvest (4, 8, and 12 weeks), gravimetric soil moisture was measured for all microcosms (Supp. Fig. S3).

In total, the experiment contained 120 microcosms. Each combination of moisture (normal, drought) × harvest time point (4, 8, 12 weeks) × plant C input (rhizodeposits, root detritus) had 10 replicate mesocosms (divided between: *n* = 6 ^13^C-labeled and *n* = 4 natural abundance controls) (see Supp. Fig. S1 for full experimental design).

### Sample collection

At each harvest, aboveground *A. barbata* biomass in planted microcosms was clipped at the base of the stems, dried at 65° C and weighed. Rhizosphere soil was collected by gently shaking the root systems to remove loosely attached soil; soil still clinging and < 2mm from roots was characterized as rhizosphere. A subset of roots + rhizosphere soil was immediately placed within a 15-mL falcon tube on dry ice and stored at −80°C. The remaining rhizosphere soil was carefully separated from the roots by hand; a subset was kept fresh at room temperature for gravimetric soil moisture and other assays (described below), and a subset was air-dried for SOC analysis. In unplanted microcosms (root detritus treatment), detritusphere soil was collected from inside the 28-μm ‘detritusphere mesh bag’; care was taken to avoid any visually apparent pieces of decaying root material. As above, subsets were stored at −80°C, fresh, and air-dried.

### SOC analysis

We isolated the mineral-associated SOC fraction on all ^13^C-labeled soils (*n* = 6) and a set of natural abundance samples from week 12 (*n* = 4), using a combined density and physical fractionation method^61^. We added 25 mL of 1.85 g cm^3^ sodium polytungstate (SPT) to 5 g of air-dried soil in a 50-mL falcon tube. To disperse aggregates, samples were shaken on a reciprocal shaker (~200 oscillations/minute) for 18 hours with glass beads. The supernatant was filtered with a glass fiber filter to separate the light particulate fraction (<1.85 g cm^−3^). The pelletized heavy fraction at the base of the tube (<1.85 g cm^−3^) was rinsed three times with deionized water to remove all excess SPT. The heavy fraction pellet was then vortexed with 25-mL deionized water and passed through a 53-μm sieve. This < 53-μm fraction (clay + fine silt) was defined as mineral-associated SOC^61,62^. Mineral-associated SOC samples were dried, ground, weighed, and analyzed for % total C and δ^13^C on an elemental analyzer coupled to an isotope ratio mass spectrometer (EA-IRMS; Costech ECS 4010, Costech Analytical Technologies, Valencia, CA, USA). Atom% ^13^C enrichment of mineral-associated SOC was calculated by subtracting atom% ^13^C of mineral-associated SOC at *t* = 0 from the atom% ^13^C in the enriched sample. The total μg of ^13^C that accumulated in the mineral-associated SOC fraction was then calculated by multiplying %C of the mineral-associated SOC fraction by the atom % 13C enrichment of the mineral-associated fraction at a given timepoint (4,8,12 weeks).

### Microbial traits

Immediately after each harvest, we measured microbial biomass C, extracellular polymeric substances (EPS), and microbial community-level growth rate and carbon-use efficiency on fresh soil. Microbial biomass C was measured via a chloroform fumigation extraction protocol on two ~3-6g soil subsamples^63^. One ~3-6g subsample was extracted in 25 mL of 0.05M K_2_SO_4_ immediately, and one ~3-6g subsample was fumigated for 7 days with chloroform prior to extraction. Both extracts were filtered (Whatman No. 40) and analyzed for total C on a Shimadzu TOC Analyzer, and for δ ^3^C on a Thermo DeltaPlus XP GasBench. The unfumigated sample was defined as dissolved organic C; microbial biomass C was calculated as the difference between the fumigated – unfumigated samples (extraction efficiency K_EC_ = 0.45). Atom% ^13^C enrichment of MBC and DOC was calculated using the same equation as described above for the mineral-associated SOC fraction.

To extract extracellular polymeric substances (EPS), we used a modified cation exchange resin extraction method^64,65^ with an added ethanol precipitation step (for detailed method, see: Sher et al.^66^). Briefly, 5-10g of soil was added to a cooled 50-mL falcon tube containing 15-mL of phosphate buffer saline and 2 g cation exchange resin (Dowex® Marathon® C, 20–50 mesh, Na^+^ form, Sigma-Aldrich, St. Louis). Samples were vortexed, shaken, and centrifuged, and 8-mL of the supernatant was passed through a 0.2 μm nylon syringe filter. EPS was precipitated from the filtrate with three, 24-mL volumes of freezer-cold, 100% ethanol and concentrated 10x. The ethanol was dried off, and the pellet was resuspended in 0.5 mL of ultrapure MilliQ water. The extract was then transferred to a pre-weighed tin capsule, dried at 70°C, and analyzed for %C and δ^13^C using the same EA-IRMS as described above. Atom% ^13^C enrichment of EPS was calculated by subtracting atom% ^13^C in natural abundance control samples from atom% ^13^C in enriched samples. The μg of ^13^C-EPS g soil^−1^ was calculated by multiplying atom% ^13^C enrichment by %C of the EPS, and dividing by the total g of soil from which EPS had been extracted^66^.

We measured community-level carbon-use efficiency and mass-specific microbial growth rates using the substrate-independent, ^18^O-H_2_O method, which captures the incorporation of isotopically labeled water into microbial DNA^67^. A 1-g sample of fresh soil was weighed into a 20-mL Wheaton glass serum vial, and a total of 50-μL of water was added to each sample – a combination of ^18^O-H_2_O (98 atom % H_2_^18^O, Isoflex, San Francisco, CA, USA) and natural abundance H_2_O – so the resulting soil water solution was ~20 atom% ^18^O. We selected this volume of water after testing for the minimum amount of water that would ensure adequate diffusion throughout the sample, while also maintaining moisture differences between normal moisture and drought treatments during the incubation period. An additional 1-g soil aliquot from a subset of 3 replicates from each moisture regime × microbial habitat combination at week 12 was incubated with 50-μL natural abundance water.

Vials were capped following the water addition. After a 72-hour incubation at room temperature, 5-mL of gas from the vial headspace was collected for CO_2_ analysis (and respiration rate) on a gas chromatograph. After sampling the headspace, the soil was removed and immediately flash frozen in liquid nitrogen and stored at −80°C. DNA was extracted from frozen soil using a Qiagen PowerSoil Pro kit and quantified using the Qubit DNA BR Assay Kit (ThermoFischer Scientific). A 50-μL aliquot of DNA was dried at 60°C in a pre-weighed silver capsule spiked with 100 μL salmon sperm DNA (to achieve the oxygen detection limit) and analyzed for δ^18^O and total O content (μg) on a Thermochemical Element Analyzer (TC/EA) coupled to an IRMS.

CUE and microbial growth rate were calculated from the amount of new DNA produced during the incubation period (tracked by ^18^O incorporation into microbial DNA during growth). The amount of new DNA produced during the incubation period (DNA_p_) is the difference in ^18^O abundance between DNA from labeled and control incubations times the proportion (by mass) of DNA that is oxygen (0.3121) divided by the length of the incubation and mass of soil incubated. The conversion mass ratios of MBC:DNA for each sample was applied to calculate mass-specific growth rate (Gr) in μg C day^−1^ g^−1^ soil:

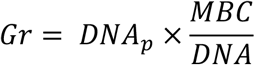

Where DNAp (μg DNA day^−1^ g^−1^ dry soil) is the DNA produced during the incubation, MBC (μg C) is microbial biomass C measured via chloroform fumigation extraction (described above), and DNA is the soil DNA concentration (μg DNA g^−1^ dry soil) determined via Qubit.

Carbon-use efficiency (CUE) was calculated as the amount of C used for growth relative to the sum of C allocated toward growth and respiration:

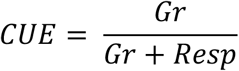

Where Resp is the average respiration rate (μg C day^−1^ g^−1^ soil) during the incubation.

We measured extracellular enzymatic potential (nmol g^−1^ soil hr^−1^) of five different enzymes related to C-, N-, and P-cycling (see Supplementary Table S8) using a fluorometric method^68^. Briefly, 1.5 g of soil, which had been stored at −80°C, was suspended in 150 mL of maleate buffer (25 mM, pH 4.95). 100 μL of soil suspension and 100 μL of substrate were pipetted into black microtiter plates, with five analytical replicates. Methylumbelliferyl (MUB) was used to calibrate all enzymes and a quench correction was performed. Plates were incubated in the dark at room temperature and fluorescence was measured every 30 min for four hours at an excitation wavelength of 365 nm and an emission wavelength of 450 nm on a BioTek Synergy HT plate reader (Bio-Tek Instruments, Winooski, VT). Enzyme activity was calculated as the increase in fluorescence over time. Cumulative exoenzyme potential (nmol g^−1^ soil hr^−1^) was calculated as the sum of average activity across all enzymes.

### Synchrotron-based scanning transmission X-ray microscopy (STXM) in combination with near edge X-ray absorption fine structure (NEXAFS) spectroscopy

STXM analysis was performed on the Molecular Environmental Sciences beamline 5.3.2.2 of the Advanced Light Source (ALS) at Lawrence Berkeley National Laboratory (California, USA)^69,70^. A single replicate of a mineral-associated SOC sample from each moisture regime × microbial habitat combination at week 12 were finely ground, suspended in Milli-Q (18.2 Ω) water and the suspension was dropcast onto Si_3_N_4_ windows (Silson Ltd.) with a micropipette and air-dried. Carbon NEXAFS spectra were collected for 3-5 regions of interest (ROIs) per treatment. Care was taken to avoid thick C areas to prevent overabsorption. Calibration was done for the C 1s edge of gaseous CO_2_ at 292.74 and 294.96 eV^71^. Spatial and spectral resolution during the measurements were 40–50 nm and 0.1 eV, respectively. Dwell time was set at 2 ms. Transmission images at energies below and at the relevant absorption edge energies were converted into optical density (OD) images [OD = ln(I_0_/I), with I_0_ being the incident photon flux and I, the transmitted flux]. Image sequences (or “stacks”) acquired at multiple energies spanning 278–330 eV for the C 1s edge were used to extract NEXAFS spectra. Clear areas of the Si_3_N_4_ membrane were used to normalize the transmission signal obtained from analyzed ROIs. All STXM data processing was done using the IDL package aXis2000^72^. C NEXAFS spectra were normalized using the Athena software package for X-ray absorption spectroscopy^73^.

### ^13^C-Nuclear Magnetic Resonance

The ^13^C chemical composition of mineral associated SOC samples were determined via solid state ^1^H-^13^C cross-polarization magic angle spinning (CP/MAS) nuclear magnetic resonance (NMR), on a 300 MHz Bruker Neo spectrometer at Lawrence Livermore National Laboratory, operating at a ^13^C Larmor frequency of 75.71 MHz. For each plant C input × moisture treatment combination (harvested at week 12), spectra were collected on a composite sample generated from 6 replicates of ^13^C-labeled samples (*n* = 6). Soil samples (1g) were packed in 4 mm ZrO_2_ rotors and spun at 10 kHz. ^13^C CP/MAS experiments were collected with a contact time of 1.5 ms and used spinal64 for proton decoupling. Each spectra was collected for 100,000-150,000 scans with a recycle delay of 2 s. α-glycine was used as a standard for setting up the CP experiment and the isotropic shift of the carboxyl groups (δ_iso_= −176.5 ppm) in α-glycine was used as the external reference for the ^13^C chemical shift scale. ^13^C spectra were deconvoluted with gaussian line shapes to determine relative fractions^74^ using the software dmfit^75^ (Supp. Fig. S8). The line widths were informed by the chemical shift regions described for each organic matter compound class: alkyl C (0-45 ppm), N-Alkyl (45-60 ppm), O Alkyl (60 – 95 ppm), Di-O-Alkyl (95-110 ppm), aryl (110-145 ppm), O-aryl (145-165 ppm), amide (165-190 ppm), and ketone (190-215 ppm)^76^. The alkyl C: O alkyl C ratio (0-45/45-110 ppm) was calculated for each sample to characterize the degree of aliphaticity, which captures the extent of microbial decomposition and transformation of organic matter^34,77^ (ratios calculated in Supp. Table S5).

### DNA extraction and quantitative stable isotope probing

DNA was extracted from rhizosphere and detritusphere soil using a Qiagen DNEasy PowerSoil Pro kit (following the manufacturer’s instructions) on three separate 0.25 g aliquots per sample, which were then combined into a single replicate. For quantitative stable isotope probing, 5 μg of DNA in 150 μL 1xTE was loaded into a 5.2 mL ultracentrifuge tube and mixed with 1.00 mL gradient buffer, and 4.60 mL CsCl stock (1.885 g mL^−1^) with a final average density of 1.725-1.730 g mL^−1^. Samples were spun in a Beckman Coulter Optima XE-90 ultracentrifuge using a VTi65.2 rotor at 176,284 RCF_avg_ at 20 °C for 108 hours. Automated sample fractionation was performed using Lawrence Livermore National Laboratory’s high-throughput SIP (‘HT-SIP’) pipeline^78^, generating 22 ~200 μL fractions per sample. The density of each fraction was measured with a Reichart AR200 digital refractometer fitted with a prism covering to facilitate measurement from 5 μL^79^. DNA was purified and concentrated using a Hamilton Microlab Star liquid handling system programmed to automate glycogen/PEG precipitations^80^. The DNA concentration of each fraction was quantified with a PicoGreen fluorescence assay (Invitrogen, Thermo Fisher)^78^.

From each mesocosm, fractions within the density range 1.650 – 1.760 g mL^−1^ and unfractionated DNA were prepared for amplicon sequencing of ITS and 16S rRNA genes. Bacterial 16S rRNA gene copies were quantified using quantitative PCR and primers 515 F and 80 6 R^81,82^. PCR conditions for 16S rRNA gene quantification were 95 °C for 2 min followed by 20 cycles of 95 °C for 30 °S, 64.5 °C for 30 °S, and 72 °C for 1 min. A total of 511 fraction libraries were sequenced for the 16S and ITS rRNA region on an Illumina MiSeq at Northern Arizona University’s Genetics Core Facility using a 300-cycle and 500-cycle v2 reagent kit, respectively. Fungal 18S rRNA gene copies were also quantified in each density fraction using primers 1380F and 1510R. PCR conditions for 18S rRNA gene quantification were 98 °C for 3 min followed by 40 cycles of 98 °C for 45 s, 60 °C for 45 s and 72 °C for 30 s. DNA fractions, and unfractionated DNA, were amplified for fungal ITS rRNA using primers ITS4F and 5.8SF. The PCR conditions for ITS amplification were 95 °C for 6 min followed by 35 cycles of 95 °C for 15 s, 55 °C for 30 s, and 72 °C for 1 min.

### Sequence processing and qSIP analysis

Paired-end 151 nt reads were filtered to remove phiX and other contaminants with bbduk v38.56 (default settings except k=31 and hdist=1) ^83^. Fastq files were then trimmed to retain nucleotides 5-140 for the 16S F/R amplicons, and 5-245/175 for the ITS F and R reads, respectively, filtered for quality (maxEE=2, truncQ=2) and used to generate amplicon sequence variants (ASVs) with DADA2 v1.20 and phyloseq v1.36 ^84,85^. Chimeric sequences were determined and removed using removeBimeraDenovo from DADA2. ASV taxonomy was determined using the RDP 16S rRNA gene database (training set 18) using RDP classifier v2.11, keeping classifications with greater than 50% confidence^86^,, and with the UNITE v8.3 database for ITS amplicons^87^. An alignment and phylogenetic tree was built using MAFFT v7.490 and FastTree v2.1.10^88,89^.

Excess atom fraction (EAF) ^13^C of bacterial and fungal DNA was quantified following the procedure described by Hungate et al. (2015)^36^ and Koch et al. (2018)^90^ with some modifications. The density of each bacterial and fungal taxon was calculated as a weighted average across the CsCl density gradient. We used qPCR-based 16S rRNA or ITS gene copy number to normalize the relative abundance of taxa in each density fraction to compute weighted average density. The difference in a taxon’s weighted average density between natural abundance (^12^C) and ^13^C-labelled soils, as well as the taxon’s GC content, was used to estimate the change in the molecular weight of DNA due to incorporation of the ^13^C isotope, expressed as EAF ^13^C. We computed estimates of EAF ^13^C for each taxon at the level of individual ultracentrifuge tubes as the difference in a taxon’s weighted average density in a ^13^C tube minus its average weighted average density across all replicate natural abundance tubes.. Preliminary analysis showed an effect of ultracentrifuge tube on estimates of weighted average density, which has previously been attributed to slight differences in the CsCl density gradients among ultracentrifuge tubes^91^. We corrected this technical error following the approach described in Morrissey et al.^91^. Estimates of EAF ^13^C were weighted by a taxon’s relative abundance in each experimental soil.

We approximated cumulative bacterial and fungal ^13^C assimilation for each sample by summing weighted EAF ^13^C values across all fungal or bacterial taxa within that sample. We estimated ‘proportional ^13^C assimilation’ for each sample – defined as the proportion of cumulative ^13^C assimilation performed by individual taxa – by dividing cumulative bacterial or fungal ^13^C assimilation by an individual taxon’s weighted EAF ^13^C. This standardized ^13^C assimilation value allowed us to compare across rhizosphere and detritusphere treatments, which received different total amounts of ^13^C. ‘Top assimilators’ for each treatment were defined by ranking families in order of their proportional ^13^C assimilation value and summing up the values of the top assimilators until we reached a threshold value of at least 50%. We characterized traits of these top assimilators by literature searches.

### Statistical analysis

All analyses were conducted in R version 4.2.1 (R Core Team, 2022). The effect of microbial habitat and moisture regime on microbial traits were analyzed with multiple regression (Supp. Table S2). In these analyses, we specifically analyzed microbial traits that did not employ the ^13^C-label (i.e., growth rate, carbon-use efficiency, total microbial biomass C, extracellular enzyme potential), to allow direct comparisons between rhizosphere and detritusphere treatments, as these treatments received different amounts of total ^13^C from rhizodeposits versus root inputs, respectively. Differences between means for microbial traits at week 12 (Fig. 1) were calculated using Tukey’s HSD (p < 0.05).

Multiple regression models were also used to analyze the effect of different microbial traits (growth rate, CUE, ^13^C-microbial biomass C, and ^13^C-EPS), moisture treatment, and time on ^13^C-mineral-associated SOC. As these models included traits that employed the ^13^C-label, they were run separately for ^13^C-rhizodeposts in the rhizosphere versus ^13^C-root litter in the detritusphere. To avoid overfitting, each trait was run in a separate model (Supp. Table S3-S4). We also analyzed relationships between these four microbial traits and ^13^C-mineral associated SOC at week 12 using simple linear regression (Fig. 2). For all models, interaction terms between factors were dropped from the model if they were not marginally significant at p < 0.1. All models were screened for normality of residuals (Shapiro-Wilk test).

Cumulative ^13^C-assimilation was compared between bacteria and fungi within each microbial habitat × moisture treatment using t-tests (Fig. 4). We used principal coordinate analysis (PCoA) of Bray-Curtis distances to visualize taxon-specific estimates of proportional ^13^C-assimilation and PERMANOVA to assess whether bacterial and fungal communities consuming ^13^C-labeled substrates differed between microbial habitats and soil moisture regimes at week 12 (Supp. Fig .S7).

## ACKNOWLEDGEMENTS

We thank Gianna Marschmann, Ella Sieradzki, Rhona Stuart, Erin Nuccio, Gareth Trubl, Cynthia Jeanette-Mancilla, Melanie Rodriguez-Fuentes, Dinesh Adhikari, Laura Adame, Peter Weber, Rachel Neurath, David Sanchez, Nameer Baker, Sarah Roy, Ilexis Chu-Jacoby, Rachel Hestrin, Emily Kline, Christina Fossum, Mengting Yuan, Alexa Nicolas, Anne Kakouridis, Donald Herman, Craig See, and Aaron Chew for assistance with soil collection, greenhouse harvests and lab analyses; Tina Winstrom at the Oxford Tract Greenhouse at UC Berkeley where the experiment was conducted, and John Bailey and Allison Smith at the Hopland Research and Extension Center where soil was collected. We thank Brad Erkkila at the Yale Analytical and Stable Isotope Laboratory and Jamie Brown at Northern Arizona University for help with stable isotope analyses; Jessica Wollard for assistance with DNA extractions, Michaela Hayer for sequencing DNA samples and conducting quantitative PC, and Christina Ramon for assistance preparing soil samples for isotope analysis. We also thank Harris Mason, April Sawvel, and Christoper Colla for assistance with ^13^C-NMR at LLNL; and Whendee Silver, Summer Ahmed, and Heather Deng for assistance with the gas chromatograph at UC Berkeley. The LLNL Soil Microbiome SFA team provided valuable feedback on experimental design and data interpretation. This research was supported by the U.S. Department of Energy, Office of Biological and Environmental Research, Genomic Science Program ‘Microbes Persist’ Scientific Focus Area (#SCW1632) at Lawrence Livermore National Laboratory (LLNL) and subcontracts to Northern Arizona University, Lawrence Berkeley National Laboratory and the University of California, Berkeley. Work conducted at LLNL was conducted under the auspices of the US Department of Energy under Contract DE-AC52-07NA27344. Work performed at Lawrence Berkeley National Laboratory was funded under U.S. Department of Energy contract number DE-AC02-05CH11231.

