## Supplementary Information for "Divergent microbial traits influence the transformation of living versus dead root inputs to soil carbon"

**Affiliations:**
[1]*Lawrence Livermore National Laboratory, Physical and Life Science Directorate, Livermore, CA, USA*
[2] *Center for Ecosystem Science and Society, Northern Arizona University, Flagstaff, AZ, USA*
[3]*University of California Berkeley, Department of Environmental Science Policy and Management, Berkeley, CA, USA*
[4] *Earth and Environmental Sciences, Lawrence Berkeley National Laboratory, Berkeley, CA, USA*
[5]*University of California Merced, Life & Environmental Sciences Department, Merced, CA, USA*

**This file contains:**
**Figures S1-S6**
**Tables S1-S7**

**Figure S1. Schematic of experimental design at a single timepoint.** Root input treatments included (1) rhizodeposits, which were tracked into the rhizosphere in microcosms planted with *Avena barbata*, and (2) root detritus, tracked in the detritusphere – defined as the mix of *A. barbata* root detritus + soil contained with a 28-µm mesh bag, buried at the center of an unplanted microcosm (see Supp. Fig. S2). Both types of root input treatment were replicated ten times ($n$ = 10) under both normal moisture (~16% soil moisture) and drought (~8% soil moisture conditions. Within each root input × moisture treatment combination, we replicated the $^{13}$C-labeled treatment 6 times ($n$ = 6, shown in red), and a natural abundance, unlabeled control 4 times ($n$ = 4, shown in blue). The same design was repeated for all three time points: 4, 8, and 12 weeks.

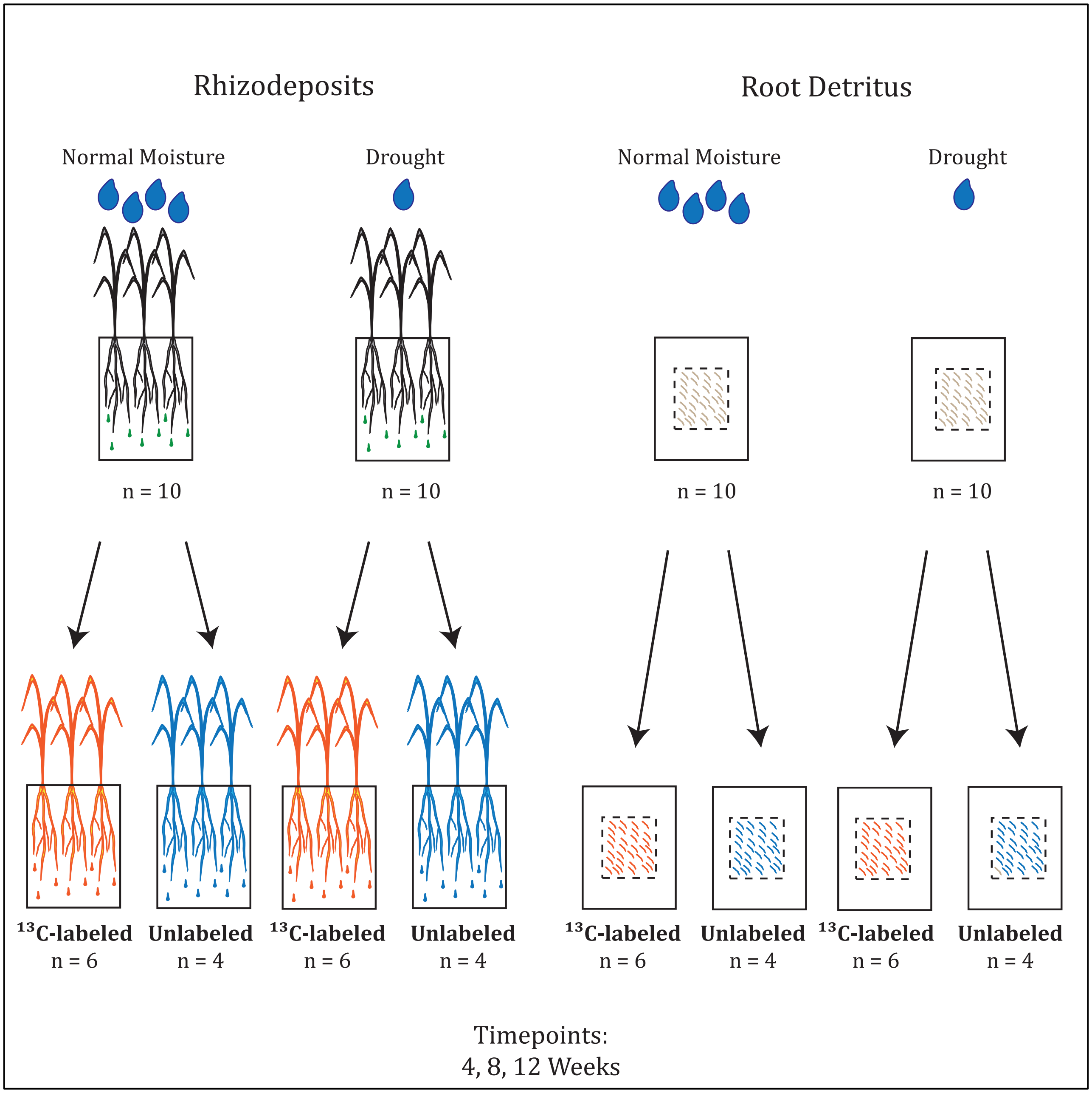

**Figure S2. Images of planted microcosms and unplanted microcosms.** (a) Planted microcosms contained living *Avena barbata* plants, whose rhizodeposits were tracked into rhizosphere soil. (b) Rhizosphere soil was defined as the soil directly clinging to roots after gently shaking them. (c) Unplanted microcosms (root detritus treatment) contained a ‘detritusphere’ 28-µm mesh bag mesh buried at the center of the microcosm, and fully covered with soil. (d) Detritusphere bag contained 1-5 mm fragments of *A. barbata* dead roots mixed with Hopland soil.

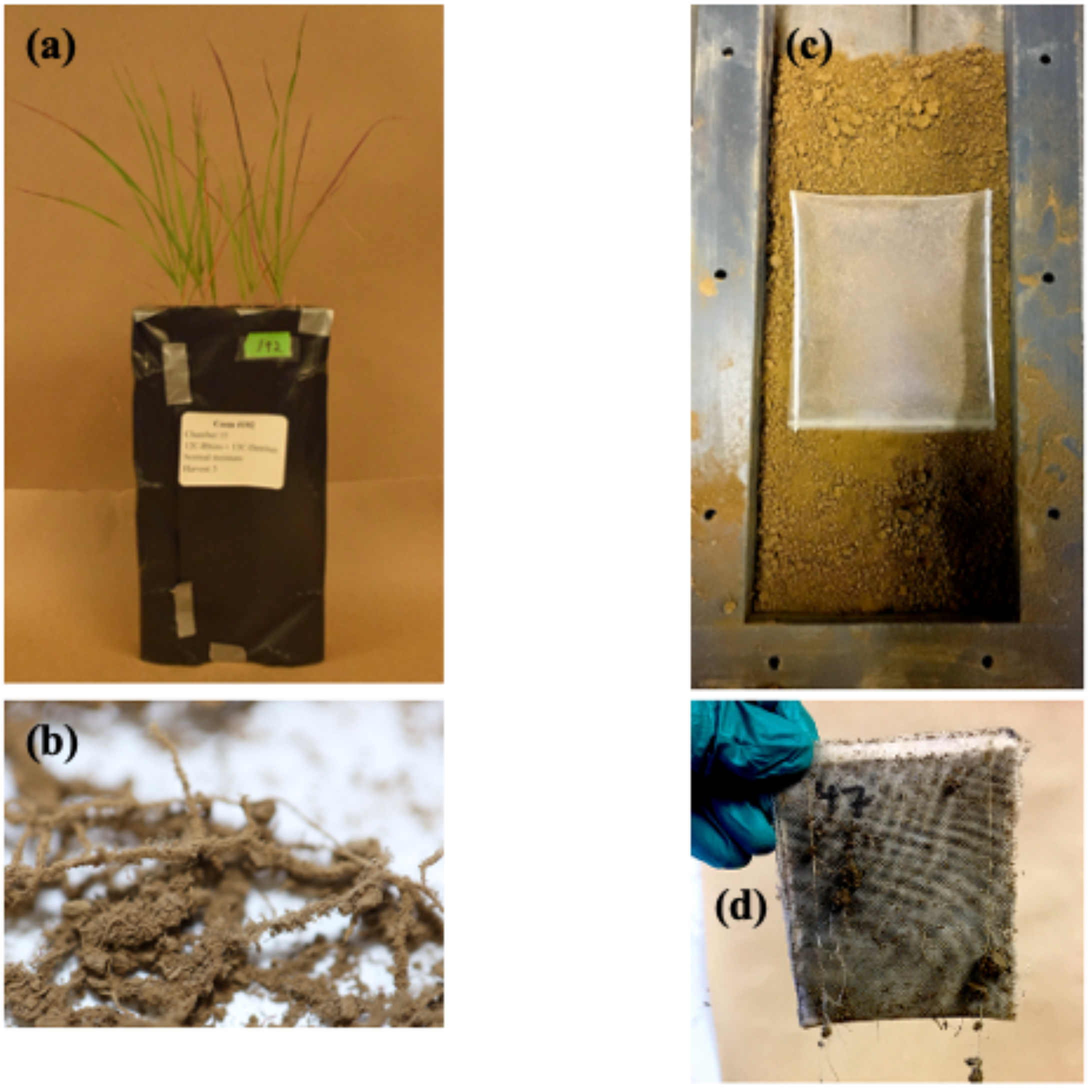

**Figure S3.** Average gravimetric soil moisture of (a) rhizosphere and (b) detritusphere soil samples at weeks 4, 8, and 12 in normal moisture (green) and drought (brown) conditions. *N* = 10 per time point. Average gravimetric moisture across time points and soil habitats was ~16% in the normal moisture treatment, and ~8% in the drought treatment. Bars indicate standard error.

**(a) Rhizosphere**

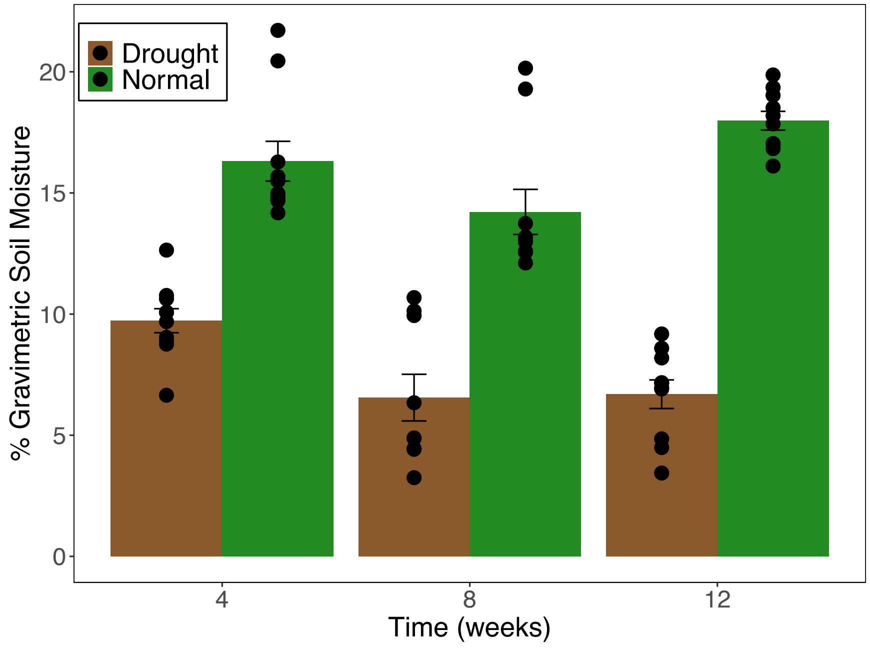

**(b) Detritusphere**

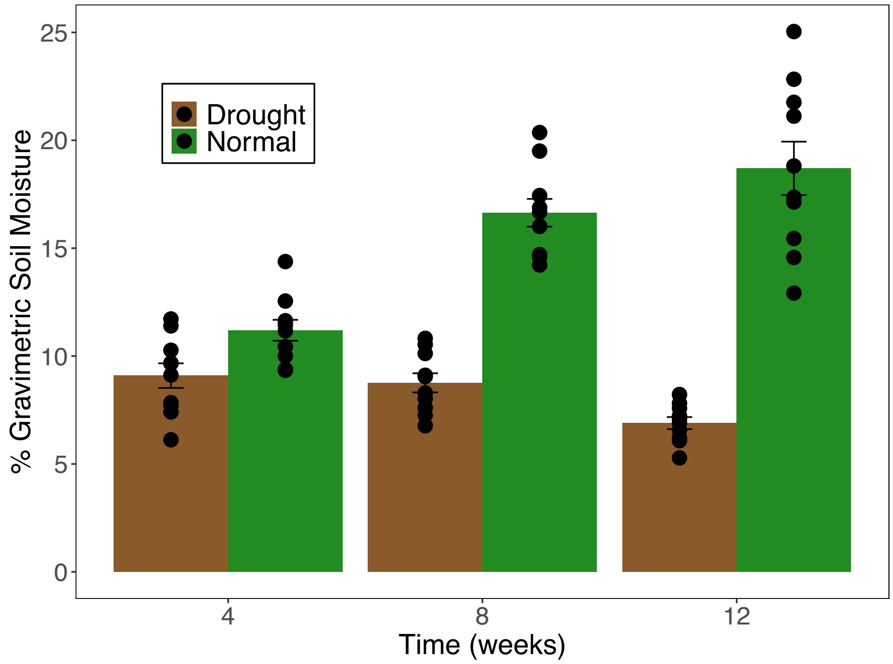

**Figure S4. Total gene copy numbers for (a) fungi (I8S) and (b) bacteria (16S) at week 12, as determined via quantitative PCR** (*n* = 4). Detritus = Detritusphere; Rhizo = rhizosphere.

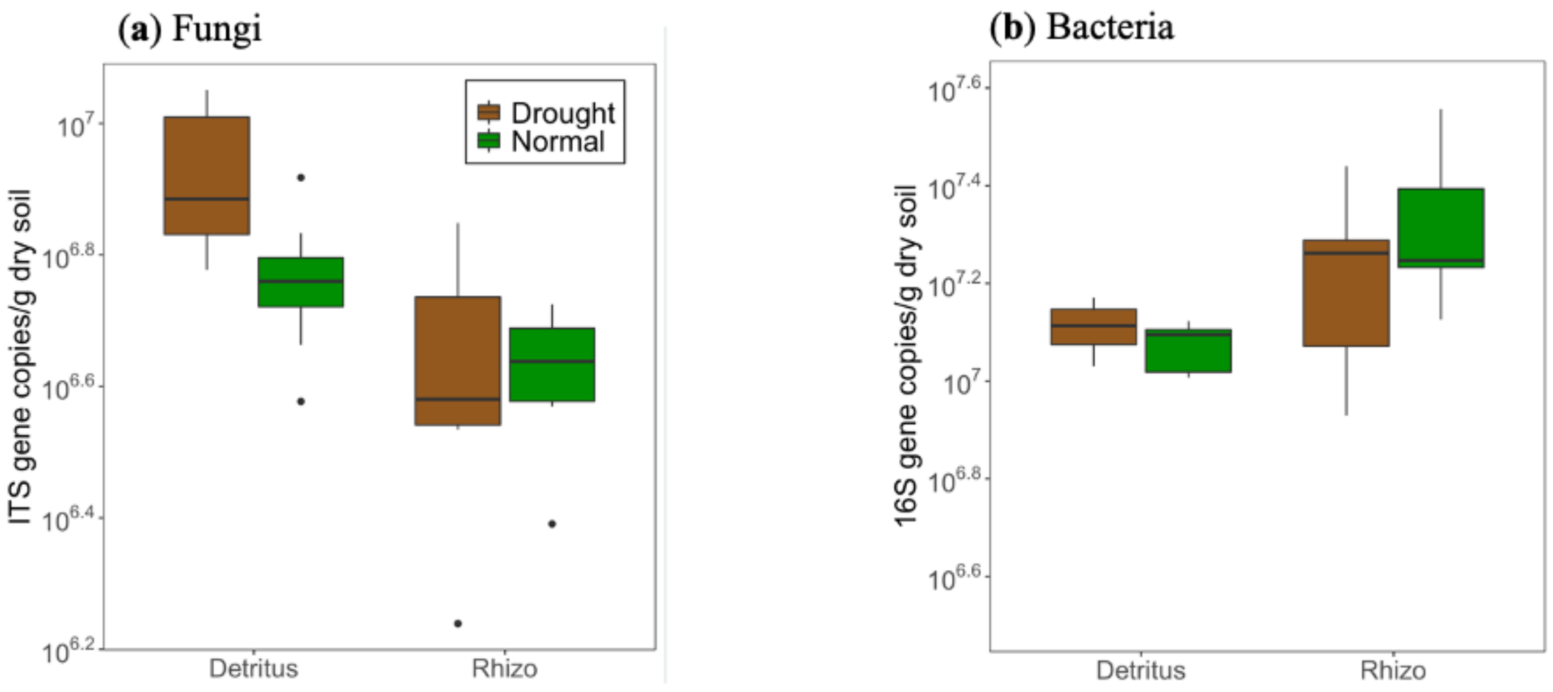

**Fig S5. Principal coordinate analysis (PCoA) of bacterial and fungal communities actively assimilating $^{13}$C-rhizodeposits in the rhizosphere and $^{13}$C-root litter in the detritusphere, based on Bray-Curtis distsances of taxon-specific proportional $^{13}$C assimilation**. Distinct bacterial (Permanova, p=0.002) and fungal (Permanova, p=0.002) communities were actively consuming $^{13}$C-labeled rhizodeposits versus $^{13}$C-labeled root litter at week 12 of the experiment. There was a marginally significant effect of soil moisture, which influenced which bacterial taxa (P=0.059) and fungal taxa (p=0.091) consumed the $^{13}$C-labeled inputs under drought.

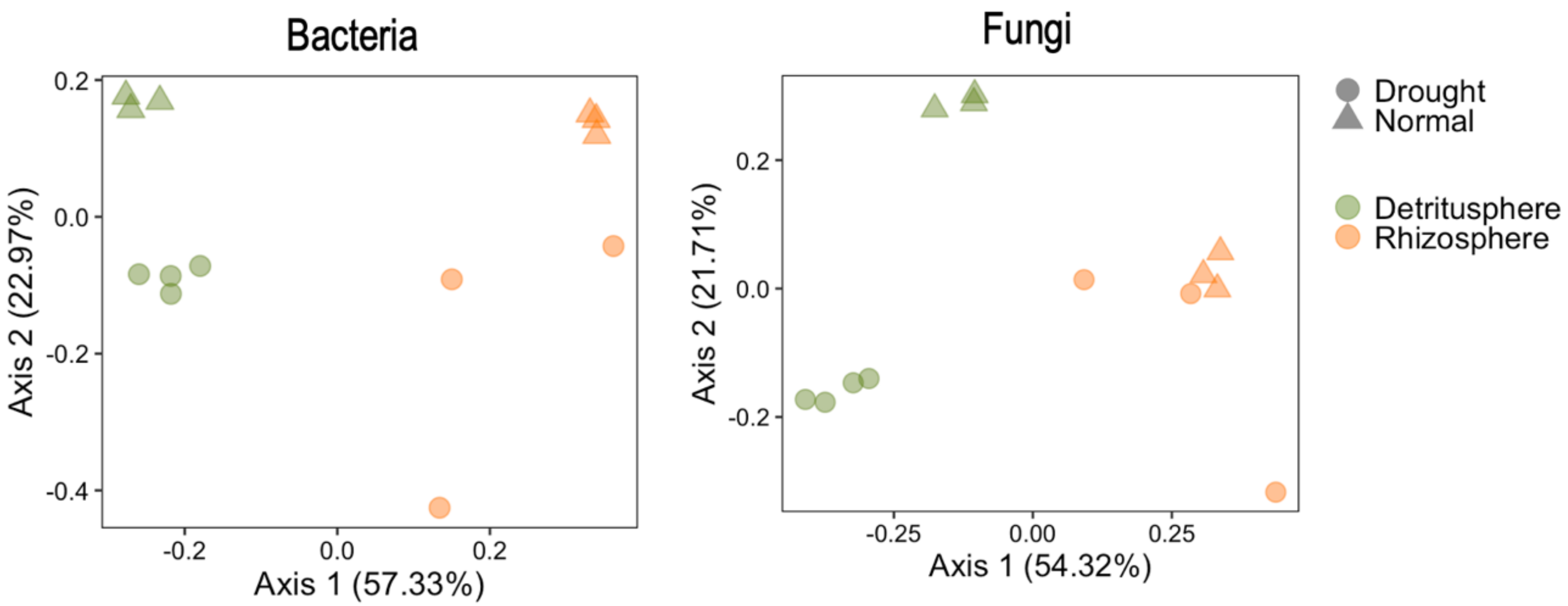

**Fig. S6. $^{13}$C CP/MAS NMR spectra of $^{13}$C-mineral associated SOC samples at week 12, shown with the corresponding deconvolution used to determine site fractions**. Samples shown under (a) normal moisture (16% soil moisture), and (b) drought conditions (8% soil moisture). Red spectra are rhizosphere samples, blue spectra are detritusphere samples. Spectra were measured on a single, composite set of 6 replicates (n = 6).

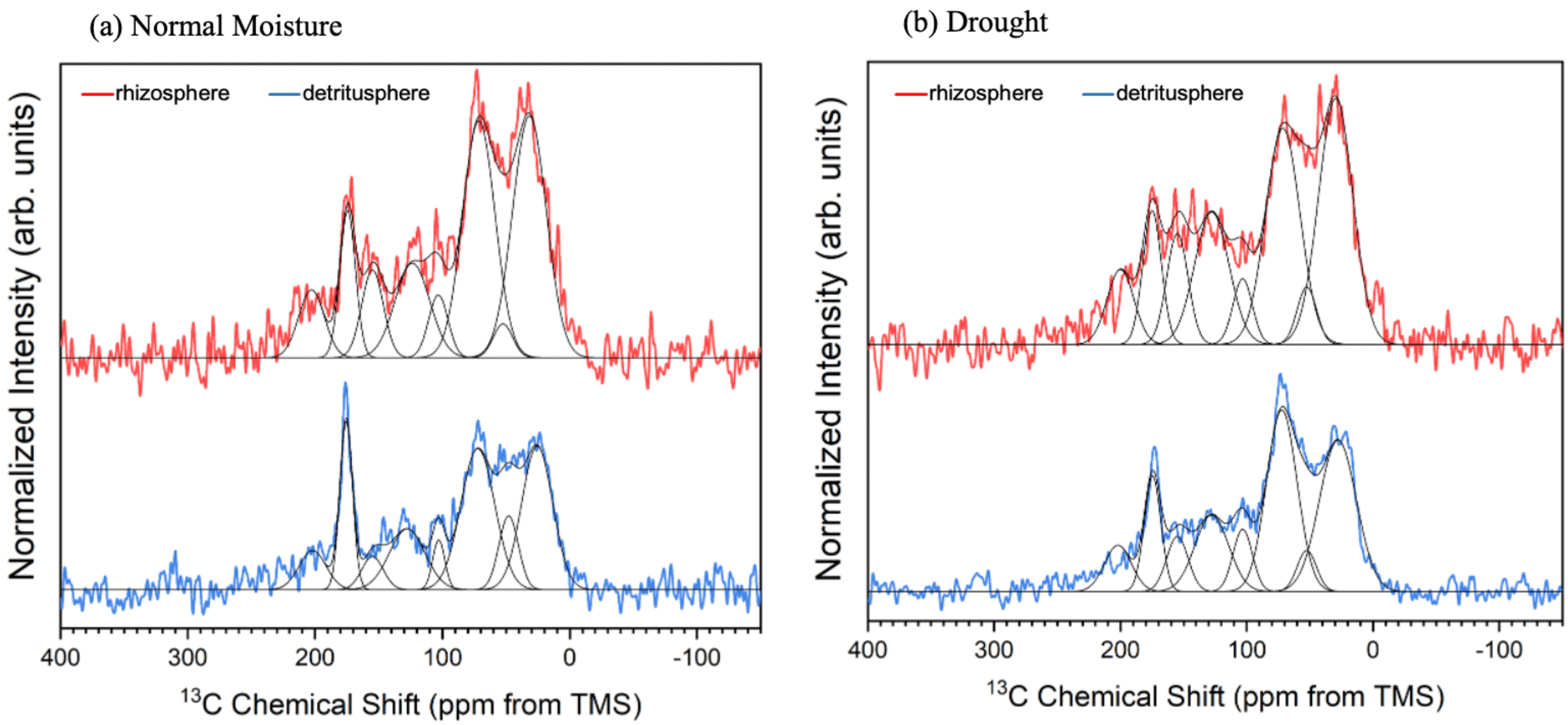

**Table S1.** Mean values of $^{13}$C-mineral associated soil organic carbon (SOC) formed over a 12-week period (4, 8, 12 weeks) from $^{13}$C-labeled (a) *Avena barbata* rhizodeposits in the rhizosphere, and (b) *A. barbata* root detritus in the detritusphere, under both normal moisture (~16% soil moisture) and drought conditions (~8% soil moisture). Values are means ± standard deviation. $N = 6$. The total amount of $^{13}$C-mineral-associated SOC formed in the rhizosphere and detritusphere are different due to different amounts of $^{13}$C-rhizodeposits versus $^{13}$C-root litter entering the soil, and thus should not be directly compared.

| **Rhizosphere $^{13}$C-mineral-associated SOC** (μg $^{13}$C g soil$^{-1}$) | | | |
|---|---|---|---|
| | 4 weeks | 8 weeks | 12 weeks |
| Normal Moisture | 25 ± 2 | 24 ± 2 | 53 ± 9 |
| Drought | 19 ± 4 | 20 ± 5 | 30 ± 3 |

| **Detritusphere $^{13}$C-mineral-associated SOC** (μg $^{13}$C g soil$^{-1}$) | | | |
|---|---|---|---|
| | 4 weeks | 8 weeks | 12 weeks |
| Normal Moisture | 69 ± 11 | 62 ± 8 | 93 ± 4 |
| Drought | 53 ± 3 | 53 ± 14 | 90 ± 27 |

**Table S2. Linear model output for microbial traits: growth rate, carbon-use efficiency, microbial biomass carbon, and cumulative extracellular enzyme activity**. Shown are linear model regression coefficients, standard errors, t and p values with factors: microbial habitat (rhizosphere, detritusphere), time (4, 8, 12 weeks), and soil moisture (drought, normal moisture). Statistical significance determined at $p < 0.05$, marginal significance determined at $p < 0.1$

| | | **Coeff.** | **Std. Error** | **t value** | **p value** |
|---|---|---|---|---|---|
| **Growth Rate** | (Intercept) | 0.004887 | 0.001335 | 3.66 | 0.000617 |
| | Rhizosphere | -0.00576 | 0.002164 | -2.661 | 0.010496 |
| | Week 8 | -0.001301 | 0.001577 | -0.825 | 0.41335 |
| | Week 12 | -0.002456 | 0.001577 | -1.557 | 0.125966 |
| | Normal Moisture | 0.001244 | 0.001278 | 0.974 | 0.335073 |
| | Rhizosphere*Week 8 | 0.003299 | 0.00305 | 1.082 | 0.284719 |
| | Rhizosphere*Week 12 | 0.007894 | 0.002393 | 3.299 | 0.001815 |
| | Rhizosphere*Normal Moisture | 0.005649 | 0.00207 | 2.729 | 0.008794 |
| **Carbon-use efficiency** | (Intercept) | 8.925 | 2.114 | 4.222 | 9.97E-05 |
| | Week 8 | 13.285 | 2.364 | 5.621 | 8.01E-07 |
| | Week 12 | -4.477 | 2.097 | -2.135 | 0.0376 |
| | Rhizosphere | 4.895 | 2.723 | 1.798 | 0.0781 |
| | Normal Moisture | 4.775 | 2.261 | 2.112 | 0.0396 |
| | Rhizosphere*Normal Moisture | 9.069 | 3.65 | 2.485 | 0.0163 |
| **Microbial Biomass C** | | | | | |
| | (Intercept) | 0.43227 | 0.04682 | 9.232 | 2.48E-14 |
| | Normal Moisture | -0.1443 | 0.05178 | -2.787 | 0.006612 |
| | Rhizosphere | -0.09132 | 0.0665 | -1.373 | 0.173401 |
| | Week 8 | 0.58822 | 0.05397 | 10.898 | < 2e-16 |
| | Week 12 | 0.06589 | 0.04942 | 1.333 | 0.186088 |
| | Rhizosphere*Normal Moisture | 0.32414 | 0.08787 | 3.689 | 0.000404 |
| **Extracellular Enzyme Activity** | (Intercept) | 2227.6 | 245.3 | 9.082 | 2.43E-08 |
| | Week 8 | -1033.3 | 394 | -2.622 | 0.01676 |
| | Week 12 | -234.2 | 285.7 | -0.82 | 0.42244 |
| | Rhizosphere | -1137.3 | 316.9 | -3.588 | 0.00196 |
| | Normal Moisture | 534.7 | 170.4 | 3.137 | 0.00542 |
| | Rhizosphere*Week 8 | 1206.6 | 490.6 | 2.46 | 0.02367 |
| | Rhizosphere*Week 12 | 731.4 | 390.9 | 1.871 | 0.07678 |

**Table S3. Linear multiple regression model output showing the effect of different microbial traits, moisture regime, and time on $^{13}C$-mineral-associated SOC in the rhizosphere.** Traits include: growth rate, carbon-use efficiency (CUE), $^{13}C$-extracellular polymeric substances (EPS), and $^{13}C$-microbial biomass C) in the rhizosphere. Shown are linear model regression coefficients, standard errors, t and p values with factors: time (4, 8, 12 weeks), and soil moisture (drought, normal moisture). Statistical significance determined at $p < 0.05$, marginal significance determined at $p < 0.1$.

| | | Coefficient | Std. Error | t value | p value |
|---|---|---|---|---|---|
| **Growth Rate** | (Intercept) | 15.294 | 2.595 | 5.893 | 1.77E-05 |
| | Growth Rate | 647.419 | 226.898 | 2.853 | 0.011 |
| | Week 8 | -4.025 | 3.991 | -1.009 | 0.3273 |
| | Week 12 | 15.363 | 2.999 | 5.122 | 8.49E-05 |
| | Normal Moisture | 11.217 | 2.926 | 3.834 | 0.00133 |
| **CUE** | (Intercept) | 11.6218 | 3.2349 | 3.593 | 0.00194 |
| | CUE | 0.2725 | 0.1521 | 1.791 | 0.08917 |
| | Week 8 | -6.2495 | 4.5657 | -1.369 | 0.18703 |
| | Week 12 | 19.5863 | 2.7877 | 7.026 | 1.09E-06 |
| | Normal Moisture | 11.2177 | 3.3182 | 3.381 | 0.00314 |
| | (Intercept) | 13.985 | 2.4376 | 5.737 | 1.08E-05 |
| **$^{13}C$-EPS** | $^{13}C$-EPS | 7.9464 | 2.8607 | 2.778 | 0.011276 |
| | Week 8 | -0.5007 | 2.9497 | -0.17 | 0.866842 |
| | Week 12 | 12.4918 | 3.6042 | 3.466 | 0.002311 |
| | Normal Moisture | 10.0975 | 2.5945 | 3.892 | 0.000841 |
| | (Intercept) | 15.938375 | 2.563884 | 6.216 | 2.42E-06 |
| **$^{13}C$-MBC** | $^{13}C$-MBC | 0.004704 | 0.027916 | 0.169 | 0.867654 |
| | Week 8 | -0.1185 | 3.886441 | -0.03 | 0.975939 |
| | Week 12 | 18.292373 | 3.375615 | 5.419 | 1.66E-05 |
| | Normal Moisture | 13.679656 | 3.080897 | 4.44 | 0.000188 |
| | (Intercept) | 15.294 | 2.595 | 5.893 | 1.77E-05 |

**Table S4. Linear multiple regression model output showing the effect of different microbial traits, moisture regime, and time on $^{13}$C-mineral-associated SOC in the detritusphere.** Traits include: growth rate, carbon-use efficiency (CUE), $^{13}$C-extracellular polymeric substances (EPS), and $^{13}$C-microbial biomass C in the detritusphere. Shown are linear model regression coefficients, standard errors, t and p values with factors: time (4, 8, 12 weeks), and soil moisture (drought, normal moisture). Statistical significance determined at $p < 0.05$, marginal significance determined at $p < 0.1$.

| | | **Coefficient** | **Std. Error** | **t value** | **P value** |
|---|---|---|---|---|---|
| **Growth Rate** | (Intercept) | 56.175 | 10.08 | 5.573 | 4.63E-06 |
| | Growth Rate | 255.471 | 1800.48 | 0.142 | 0.888115 |
| | Week 8 | -4.433 | 6.267 | -0.707 | 0.484828 |
| | Week 12 | 30.492 | 7.303 | 4.175 | 0.000235 |
| | Normal Moisture | 8.784 | 5.216 | 1.684 | 0.102578 |
| **CUE** | (Intercept) | 55.319 | 7.2964 | 7.582 | 1.87E-08 |
| | CUE | 0.2226 | 0.5708 | 0.39 | 0.699 |
| | Week 8 | -7.6687 | 9.4385 | -0.812 | 0.423 |
| | Week 12 | 31.1428 | 6.6621 | 4.675 | 5.83E-05 |
| | Normal Moisture | 8.0453 | 5.4248 | 1.483 | 0.148 |
| **$^{13}$C-EPS** | (Intercept) | 51.3335 | 13.3971 | 3.832 | 0.000605 |
| | $^{13}$C-EPS | 0.7315 | 1.8221 | 0.401 | 0.690916 |
| | Week 8 | -3.9669 | 5.6851 | -0.698 | 0.4907 |
| | Week 12 | 32.83 | 7.2294 | 4.541 | 8.48E-05 |
| | Normal Moisture | 11.6908 | 6.44 | 1.815 | 0.079483 |
| **$^{13}$C-MBC** | (Intercept) | 60.52889 | 8.72309 | 6.939 | 1.52E-07 |
| | $^{13}$C-MBC | -0.03747 | 0.06734 | -0.556 | 0.582 |
| | Week 8 | -2.8095 | 7.93874 | -0.354 | 0.726 |
| | Week 12 | 29.55011 | 6.16577 | 4.793 | 4.89E-05 |
| | Normal Moisture | 9.23236 | 5.44379 | 1.696 | 0.101 |

**Table S5. $^{13}$C-NMR assigned organic matter compounds classes for peak regions.** The above table (a) shows the compound classes assigned for each peak region. The bottom table (b) shows the % signal intensity in composite samples (n = 6) of mineral-associated SOC for each treatment at Week 12. Alkyl/O-alkyl ratio (A/OA) ratio is calculated by dividing % signal intensity in the 0-45 ppm region by the 45-110 ppm region.

(a)

| Compound Class | Expected Peak Regions (ppm) | Center | Width |
|---|---|---|---|
| Alkyl | 0 – 45 | 22.5 | 45 |
| N-Alkyl/Methoxyl | 45 – 60 | 52.5 | 15 |
| O-Alkyl | 60 – 95 | 77.5 | 35 |
| Di-O-Alkyl | 95 – 110 | 102.5 | 15 |
| Aryl | 110 – 145 | 127.5 | 35 |
| O-Aryl | 145 – 165 | 155 | 20 |
| Amide/Carboxyl | 165 – 190 | 177.5 | 25 |
| Ketone | 190 – 215 | 202.5 | 25 |

(b)

| Treatment | Compound Class | Position | Width | % Signal Intensity | A/OA Ratio |
|---|---|---|---|---|---|
| Detritusphere - Normal Moisture | Alkyl | 25.65 | 28.66 | 25.65 | 0.65 |
| | N-Alkyl/Methoxyl | 48.07 | 18 | 8.28 | |
| | O-Alkyl | 72.28 | 31.36 | 27.56 | |
| | Di-O-Alkyl | 103 | 11.22 | 3.48 | |
| | Aryl | 127.75 | 32.88 | 12.46 | |
| | O-Aryl | 155 | 20 | 4.13 | |
| | Amide/Carboxyl | 175.66 | 11.79 | 12.33 | |
| | Ketone | 202 | 25 | 6.1 | |
| Detritusphere - Drought | Alkyl | 27.91 | 34.5 | 27.89 | 0.74 |
| | N-Alkyl/Methoxyl | 52.5 | 18 | 3.95 | |
| | O-Alkyl | 72.32 | 28.5 | 27.63 | |
| | Di-O-Alkyl | 103.3 | 18 | 5.98 | |
| | Aryl | 127.75 | 31.38 | 12.84 | |
| | O-Aryl | 155 | 20 | 5.88 | |
| | Amide/Carboxyl | 174.64 | 15.64 | 9.63 | |
| | Ketone | 202 | 25 | 6.18 | |
| Rhizosphere Drought | Alkyl | 29.64 | 32 | 28.48 | 0.86 |
| | N-Alkyl/Methoxyl | 52.5 | 18 | 3.74 | |
| | O-Alkyl | 71.49 | 32 | 25.01 | |

| | | | | | |
|---|---|---|---|---|---|
| | Di-O-Alkyl | 103.3 | 18 | 4.27 | |
| | Aryl | 127.75 | 32 | 15.32 | |
| | O-Aryl | 155 | 20 | 8.02 | |
| | Amide/Carboxyl | 174.99 | 17.34 | 8.38 | |
| | Ketone | 200 | 25 | 6.78 | |
| Rhizosphere Normal Moisture | Alkyl | 31.75 | 31.38 | 30.38 | 0.86 |
| | N-Alkyl/Methoxyl | 52.5 | 18 | 2.45 | |
| | O-Alkyl | 71.49 | 30 | 28.46 | |
| | Di-O-Alkyl | 103.3 | 18 | 4.52 | |
| | Aryl | 123.69 | 32 | 12.13 | |
| | O-Aryl | 155 | 20 | 7.03 | |
| | Amide/Carboxyl | 174.54 | 14.57 | 8.57 | |
| | Ketone | 202.93 | 23.76 | 6.46 | |

**Table S6. Bacterial and fungal taxa top $^{13}C$ assimilators, accounting for the majority of cumulative $^{13}C$ assimilation.** Proportional $^{13}C$ assimilation values indicate the proportion of cumulative $^{13}C$ assimilation for a given taxa. Shown below are the amplicon sequence variants (ASVs) which accounted for >50% of proportional $^{13}C$ assimilation, and were defined as 'top assimilators' (see Methods section). $N = 4$.

*(a) Rhizosphere – Normal Moisture*

| | **Phylum** | **Family** | **Genus** | **Proportional Assimilation** |
|---|---|---|---|---|
| **Bacteria** | Firmicutes | Bacillaceae | Unclassified | 0.175 |
| | Proteobacteria | Comamonadaceae | Rhizobacter | 0.068 |
| | Firmicutes | Bacillaceae | Neobacillus | 0.041 |
| | Firmicutes | Bacillaceae | Unclassified | 0.034 |
| | Firmicutes | Bacillaceae | Neobacillus | 0.033 |
| | Proteobacteria | Bradyrhizobiaceae | Bradyrhizobium | 0.027 |
| | Proteobacteria | Sphingomonadaceae | Sphingomonas | 0.022 |
| | Bacteroidetes | Chitinophagaceae | Niastella | 0.021 |
| | Proteobacteria | Burkholderiaceae | Paraburkholderia | 0.018 |
| | Actinobacteria | Streptomycetaceae | Streptacidiphilus | 0.018 |
| | Acidobacteria | Gp3 | Unclassified | 0.017 |
| | Verrucomicrobia | Spartobacteria | Unclassified | 0.015 |
| | Proteobacteria | Rhizobiaceae | Rhizobium | 0.014 |
| **Fungi** | Ascomycota | Aspergillaceae | Penicillium | 0.166 |
| | Basidiomycota | Piskurozymaceae | Solicoccozyma | 0.104 |
| | Basidiomycota | Hoehnelomycetaceae | Atractiella | 0.086 |
| | Basidiomycota | Thelephoraceae | Tomentella | 0.055 |
| | Mucoromycota | Rhizopodaceae | Rhizopus | 0.047 |
| | Ascomycota | Pleosporaceae | Bipolaris | 0.045 |

*(b) Rhizosphere – Drought*

| | **Phylum** | **Family** | **Genus** | **Proportional Assimilation** |
|---|---|---|---|---|
| **Bacteria** | Firmicutes | Bacillaceae 1 | Unclassified | 0.162 |
| | Firmicutes | Bacillaceae 1 | Neobacillus | 0.085 |
| | Firmicutes | Bacillaceae 1 | Neobacillus | 0.067 |
| | Actinobacteria | Streptomycetaceae | Streptacidiphilus | 0.064 |
| | Firmicutes | Bacillaceae 1 | Unclassified | 0.031 |
| | Actinobacteria | Streptomycetaceae | Streptomyces | 0.028 |
| | Actinobacteria | Streptomycetaceae | Streptomyces | 0.020 |
| | Acidobacteria | Gp3 | Unclassified | 0.020 |
| | Proteobacteria | Bradyrhizobiaceae | Bradyrhizobium | 0.018 |

| | | | | |
|---|---|---|---|---|
| | Proteobacteria | Sphingomonadaceae | Sphingomonas | 0.015 |
| **Fungi** | Ascomycota | Aspergillaceae | Penicillium | 0.222 |
| | Ascomycota | Ceratostomataceae | Unclassified | 0.169 |
| | Basidiomycota | Piskurozymaceae | Solicoccozyma | 0.058 |
| | Ascomycota | Pleosporaceae | Bipolaris | 0.047 |

*(c) Detritusphere – Normal Moisture*

| | **Phylum** | **Family** | **Genus** | **Proportional Assimilation** |
|---|---|---|---|---|
| **Bacteria** | Actinobacteria | Streptomycetaceae | Streptacidiphilus | 0.321 |
| | Proteobacteria | Burkholderiaceae | Paraburkholderia | 0.040 |
| | Bacteroidetes | Chitinophagaceae | Niastella | 0.039 |
| | Firmicutes | Bacillaceae | Bacillus | 0.032 |
| | Firmicutes | Bacillaceae | Neobacillus | 0.029 |
| | Actinobacteria | Streptomycetaceae | Streptacidiphilus | 0.022 |
| | Firmicutes | Bacillaceae | Neobacillus | 0.021 |
| **Fungi** | Ascomycota | Lasiosphaeriaceae | Echria | 0.226 |
| | Ascomycota | Ceratostomataceae | Unclassified | 0.148 |
| | Ascomycota | Lasiosphaeriaceae | Echria | 0.061 |
| | Ascomycota | Aspergillaceae | Penicillium | 0.058 |
| | Ascomycota | Pleosporaceae | Bipolaris | 0.052 |

(d) *Detritusphere – Drought*

| | **Phylum** | **Family** | **Genus** | **Proportional Assimilation** |
|---|---|---|---|---|
| **Bacteria** | Actinobacteria | Streptomycetaceae | Streptacidiphilus | 0.230 |
| | Actinobacteria | Streptomycetaceae | Streptomyces | 0.062 |
| | Firmicutes | Bacillaceae | Bacillus | 0.050 |
| | Proteobacteria | Burkholderiaceae | Paraburkholderia | 0.036 |
| | Firmicutes | Bacillaceae | Neobacillus | 0.036 |
| | Bacteroidetes | Chitinophagaceae | Niastella | 0.026 |
| | Firmicutes | Bacillaceae | Neobacillus | 0.024 |
| | Actinobacteria | Streptomycetaceae | Streptacidiphilus | 0.024 |
| | Proteobacteria | Rhizobiaceae | Agrobacterium | 0.022 |
| **Fungi** | Ascomycota | Ceratostomataceae | Unclassified | 0.349 |
| | Ascomycota | Aspergillaceae | Penicillium | 0.091 |
| | Ascomycota | Lasiosphaeriaceae | Echria | 0.081 |

**Table S7. List of extracellular enzymes assayed.** Table includes enzyme names, functions and substrates used in assay.

| Enzyme | Enzyme function | Substrate |
|---|---|---|
| Exoglucanase | Releases cellobioside from cellulose | 4-Methylumbelliferyl β-D-cellobioside |
| β-Glucosidase | Releases glucose from cellulose and other B-glucans | 4- Methylumbelliferyl β-D-glucopyranoside |
| β-Xylosidase | Degrades hemicellulose | 4- Methylumbelliferyl β-D-xylopyranoside |
| Exochitinase (N-acetyl-β-glucosaminidase) | Releases glucosamine from chitin | 4- Methylumbelliferyl N-acetyl-β-D-glucosaminide |
| Acid Phosphatase | Releases inorganic phosphorus from organic phosphorus | 4-Methylumbelliferyl phosphate |
